# Automated Synthesis of Wireframe DNA Nanotubes

**DOI:** 10.1101/2024.01.18.576297

**Authors:** Patricia Islas, Casey M. Platnich, Yasser Gidi, Ryan Karimi, Lorianne Ginot, Gonzalo Cosa, Hanadi F. Sleiman

**Affiliations:** Department of Chemistry, McGill University, 801 Sherbrooke Street West, Montreal, Quebec H3A 0B8, Canada

## Abstract

DNA nanotechnology has revolutionized our ability to position matter at the nanoscale, but the preparation of DNA-based architectures remains laborious. To facilitate the formation of custom structures, we report a fully automated method to produce sequence- and size-defined DNA nanotubes. By programming the sequential addition of desired building blocks, rigid DX-tile-based DNA nanotubes (and flexible wireframe DNA structures) are attained, where the total number of possible constructs increases as a power function of the number of different units available. Using single-molecule fluorescence imaging, the kinetics and yield of each synthetic step can be quantitatively determined, revealing differences in self-assembly dynamics as the nanotube is built up from the solid support and providing new insights into DNA self-assembly. The exploitation of automation for both assembly and analysis (tthrough an *ad-hoc* developed K-means clustering algorithm) facilitates a workflow wherein the synthesis parameters may be iteratively improved upon, demonstrating how a single-molecule ‘assembly-analysis-optimization’ sequence can be used to generate complex, non-covalent materials in good yield. The presented synthetic strategy is generalizable, making use of equipment already available in most standard laboratories and represents the first fully automated supramolecular assembly on a solid support.

## INTRODUCTION

In the last century, automated synthetic methods revolutionized the preparation of sequence-defined covalent polymers such as peptides, DNA, and RNA.^1, 2^ More recently, automated preparations for polysaccharides^3, 4^ and other covalent structures^5, 6^ have been reported. These methodologies rely on the time-programmed, sequential addition of functionally protected monomers (amino acids, nucleotides, and saccharides, respectively), rendering polymers with defined composition and structure. Automation has also empowered the exploration of chemical space in the synthesis of small molecules^7, 8^ from predefined building blocks, expanding the associated functionality of these species.^9, 10^ For supramolecular constructs, much like for their covalent congeners, the number and arrangement of constituent units dictate their form and function. Controlling the position of building blocks confers these assemblies with dynamic and mechanical properties that can be harnessed towards applications in nanoscale positioning, sensing, and triggered delivery.^11–13^ Automating the synthesis and characterization of supramolecular constructs thus promises to advance nanotechnology through the rapid preparation of new constructs and the streamlined evaluation of different subcomponent arrangements. To date, however, this automation has remained elusive in supramolecular chemistry. While higher-order structures based on non-covalent interactions are increasingly reported in the fields of materials engineering and biosensing due to their dynamic properties,^11, 14^ these materials are typically prepared manually, with synthesis, structural characterization, and analysis steps separated temporally and spatially.

DNA is a fully programable material with well-defined properties and assembly motifs,^15^ making it an ideal candidate to explore the automated synthesis of supramolecular architectures. Utilizing DNA tile assembly^16, 17^ and DNA origami methods,^18^ scientists have used DNA extensively to create increasingly intricate and functionally diverse^19–21^ synthetic structures at the nanoscale.^22–24^ Origami assembly, however, requires hundreds of unique DNA sequences for self-assembly to occur, and conventional tile assembly methods typically creates periodic structures with little ability to selectively place functional groups. Working with wireframe DNA structures characterized by a minimal number of DNA sequences and full addressability - consisting of DNA-based ‘rungs’ and ‘linkers’ - we previously reported a manual, surface-grafted synthesis for DNA nanotubes, marking the first step toward a fully automated supramolecular assembly platform.^25^ Our method, however, required tens of hours of manual assembly, partly as the necessary chemical information required to optimize the synthesis steps, e.g. kinetics of hybridization, was lacking . Furthermore, this previous method comprised DNA and hybrid organic-DNA components bearing synthetic aromatic units that required extensive purification and involved a manual post synthetic analysis, rendering the assembly and characterization time-intensive and laborious.

Here, we report a versatile, fully automated method to synthesize and directly characterize surface-grafted supramolecular constructs (Fig. 1). Two different DNA-based designs with different morphologies and flexibilities were chosen to test the versatility of the methodology. The main design is derived from a rigid DX-tile-wireframe hybrid nanotube (DxNT) recently reported by our group,^26^ while a second, highly flexible wireframe structure^25^ was also prepared. Combining single-molecule kinetics,^27^ microfluidics, and structural design (to both minimize labor in building block preparation and to enable kinetic studies), we retrieved the rates of building block addition for different nanotube lengths to inform and optimize the duration of individual steps in the automated synthesis. Exploiting single-molecule photobleaching experiments with a newly developed algorithm based on K-means clustering for automated data screening enabled the real-time assessment of structure and yield of as-prepared DNA nanomaterials, allowing us to rapidly test assembly protocols, analyze the outcomes, and iteratively improve our procedures. Combining these individual elements, we streamline a fully automated construction strategy of supramolecular structures, relying exclusively on self-assembly of DNA building blocks at room temperature. Importantly, the use of automation allows the order of different subunits to be easily tuned, providing a facile way to construct a myriad of different structures based on differing building block positioning.

**Figure 1.**
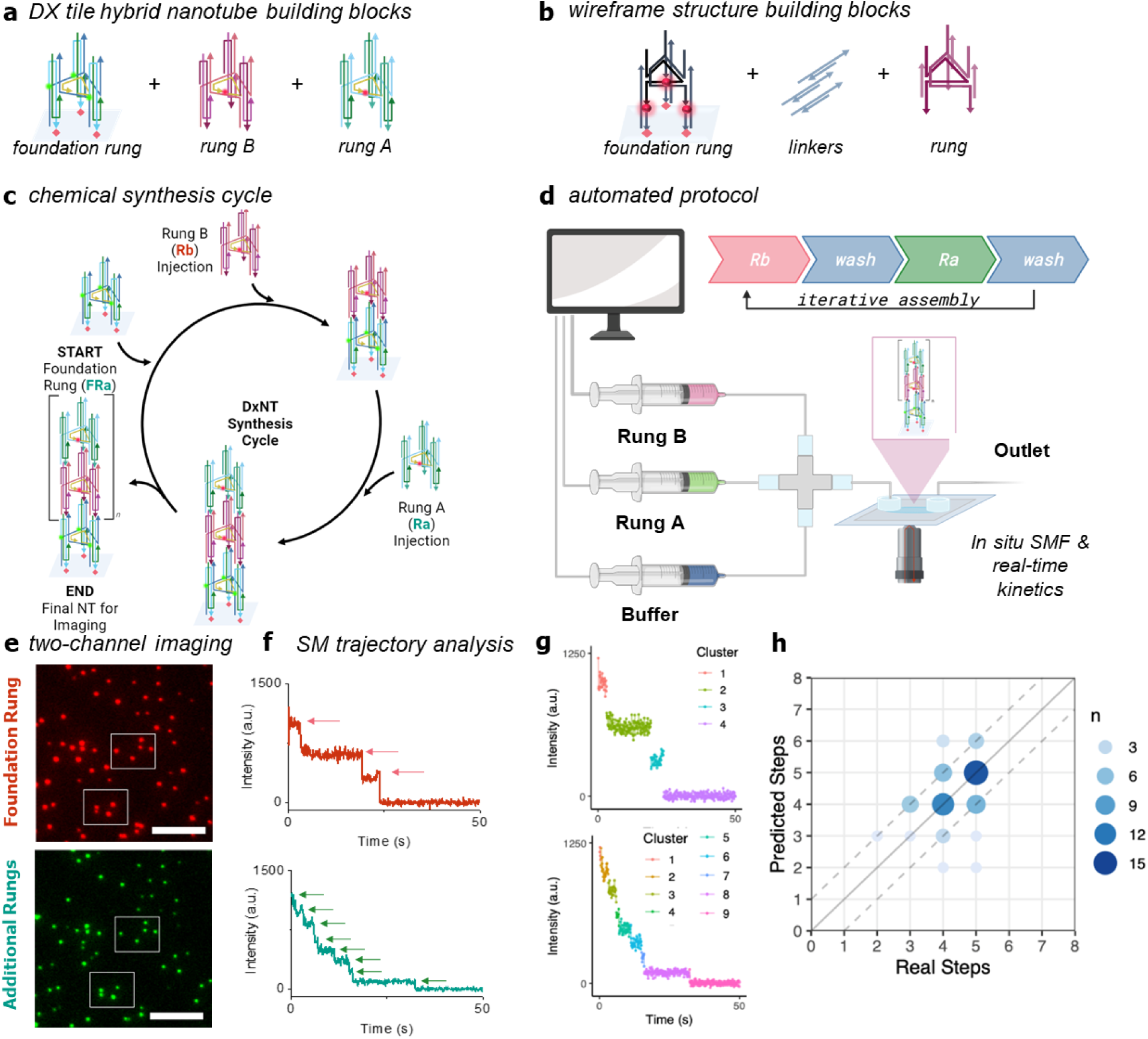
Synthesis protocols were followed for two structures, **a**. a DX tile – wireframe hybrid DNA nanotube design (DxNT) where each cycle adds two types of rungs (A and B); and **b.** a flexible DNA wireframe design, adding a set of three linkers and one rung per cycle. Summary of automated synthesis protocols. **c.** Synthesis cycle to construct a DX tile – wireframe DNA nanotube design (DxNT) one unit at a time. The foundation rung is attached to the surface; rungs are sequentially added during each cycle. The number of cycles dictates the final size and length of the structure. **d.** Schematic of prototype DNA nanotube synthesizer. Rungs (A and B) for DxNT, or rungs and linkers for the flexible DNA wireframe nanotube and buffer, are automatically pumped onto the glass coverslip in a time-programmed sequence, facilitating control over length and sequence in the final assembly. In situ analysis is performed by single-molecule fluorescence microscopy as swhon in pales **e-h. e** Sample images showing colocalization of the foundation rung (red, ATTO647N) and the subsequent rungs (green, Cy3). Scale bars = 10 μm. **f.** Photobleaching the nanotubes one color at a time allows for the stoichiometry of the final assembly to be assessed by counting the number of photobleaching steps in each trajectory. **g.** Automated clustering of intensities to extract the number of photobleaching steps enables rapid screening for synthetic success. **h.** Predicted steps, generated using our automated clustering algorithm, correlate strongly with the real steps (counted manually) for a nanotube with five Cy3-labelled rungs; dashed lines indicate population within an error of 1 count (92%).

Forming our structures directly on a passivated glass coverslip using syringe pumps^28^ facilitated the *in situ* imaging of the resulting constructs using single-molecule fluorescence microscopy, supporting their on-chip use as biophysical probes and biosensors (where we hypothesize that by incorporating aptamers and/or proteins, the nanotubes can be transformed into a multiplexing sensor).^29–31^ Surface immobilization, in conjunction with our flow setup, also allowed for the assessment of structure quality and yield at each step,^25, 32, 33^ providing a mechanistic understanding of DNA self-assembly. Incorporation of a distinct rung - bearing toeholds at the junction with the lower unit - at different levels of the nanotube assembly enabled in turn detaching the nanotube from the surface by using the strand displacement strategy.^27, 34^

While we have focused on the production of elongated DNA nanotubes built from single, rigid, tile-based units, our methodology may be used to build other modular DNA nanostructures, as demonstrated by the formation of flexible wireframe structures. The method should be further extendable to encompass other motifs, including hybrid protein-DNA structures. Using a minimal number of DNA strands and simple equipment, the approach presented herein is adaptable, user-friendly, and integrates both synthesis and characterization, paving the way for a new generation of custom, self-assembled structures.

## RESULTS AND DISCUSSION

### Building blocks and experimental design

The DNA nanotubes were made up of complementary alternating units, prepared fully from DNA following sequence planning. Our primary design, DX-NTs, carried only one type of building block – “rungs” – with two subtypes, namely A and B, with complementary sticky ends. These rungs incorporate double-crossover (DX) tile-like DNA edges to achieve structural rigidity and bear an h-motif structure permitting the assembly around a core strand into desired rung units (Fig. 1a). The “h-strand” provides two complementary 5-nucleotide sticky ends at each of its two termini, to bind to a lower (5’end) and upper (3’ end) rung. As such, three identical h-strands are incorporated in the structure providing three 5-nucloetide sticky ends above and three below. Three additional sticky ends above and three below (for a total of six in each side of the rung) are provided by the two inter-locked strands - T1 (green) and T2 (blue), see Fig. 1a - used in each of the three columns of the rung, where each provide a 5-nucleotide sticky end (above and below, respectively). A central strand completes this rung structure. Synthesis of the rung units in DxNTs thus required the annealing of four different commercially available DNA strands.^26^ Flexible wireframe DNA structures in turn required “rungs” and “linkers” where the self-assembly of the rung units occurs upon annealing six different commercially available DNA strands. Linkers consist of double stranded DNA segments that are also annealed *a priori*.^25, 27, 35^ Here, due to the relatively short persistence length of double-stranded DNA and the presence of nicks along the structure, flexible constructs are formed. In all cases, structures were characterized by polyacrylamide gel electrophoresis.

To initiate the synthesis of DxNTs, “foundation rungs”, bearing one biotin at each of their three pillars but otherwise identical in DNA sequence to rung A, (Fig. 1a) were first immobilized on the coverslip surface. These foundation rungs contained three biotin anchors to trivalently bind the construct at random positions on a PEG-passivated^28^ and streptavidin-decorated surface. Each foundation rung additionally contained three fluorophores (either Cy3 or ATTO647N), for structural characterization (see below). Likewise for flexible wireframe structures, their foundation rungs - formed from nine DNA strands, see Fig. 1b – also carried three biotin anchors and three fluorescent tags (see Fig. S1-2 for polyacrylamide gel electrophoresis).

To assemble the nanotubes, individual components (rungs A and B for DxNTs, rungs and linkers for flexible structures) were next delivered in a time-programmed sequence (Fig. 1c) using our home-built “synthesizer”, see Fig. 1d, consisting of standard laboratory syringe pumps controlled using a custom Python algorithm and connected to our glass coverslips using microfluidic tubing and silicone ports (Supporting Information, Section 4.1). Upon injection of the desired building block in buffer solution, an incubation period allowed each successive unit to hybridize. Using our automated setup, critical parameters for hybridization, including flow rate, flow time, incubation time, and volume delivered can be programmed, facilitating the fine-tuning of all synthetic parameters with minimal time input.

### Mechanistic studies and automation optimization

To gain a mechanistic understanding of the self-assembly of surface-grafted DNA nanostructures and to optimize their automated assembly, we subsequently monitored their construction in real-time upon delivery of building blocks (Fig. 2a, also Fig. S4). To this end, we utilized a total internal reflection fluorescence (TIRF) microscope suited with an EMCCD camera and a range of excitation wavelengths and emission filters to enable selectively exciting and monitoring single yellow (Cy3) and red (Cy5 or ATTO647N) emissive fluorophores real-time, at 10 frames/s sampling rate (Fig. S3). Upon addition of Cy3 labelled foundation rungs (three dyes per rung) and using our syringe pump system, we next constructed in a stepwise fashion desired assemblies incorporating unlabeled rungs (0, 3, 8, 13, 18 and 28 for DxNT). Next, we flowed in a Cy5 fluorescently-labelled subunit in the presence of photostabilizers (required to improve the photon budget of fluorophores, see Supporting Information, Section 7).^36, 37^ Exploiting our automated assembly method with real-time single particle imaging capabilities, we observed real-time the hybridization of the 2^nd^, 5^th^, 10^th^, 15^th^, 20^th^, and 30^th^ level for the DxNT (Fig. 2b, all fittings in Supporting Information, Section 7.2). A similar experimental design enabled monitoring the arrival time of the 2^nd^, 6^th^, and 11^th^ rung and linker for the flexible wireframe structure (Fig. 2c). Average binding times were determined by summation and subsequent fitting of single addition events recorded in fluorescence intensity trajectories from individual nanotubes monitored in parallel over time in a field of view (∼ 100 nanotubes).

**Figure 2.**
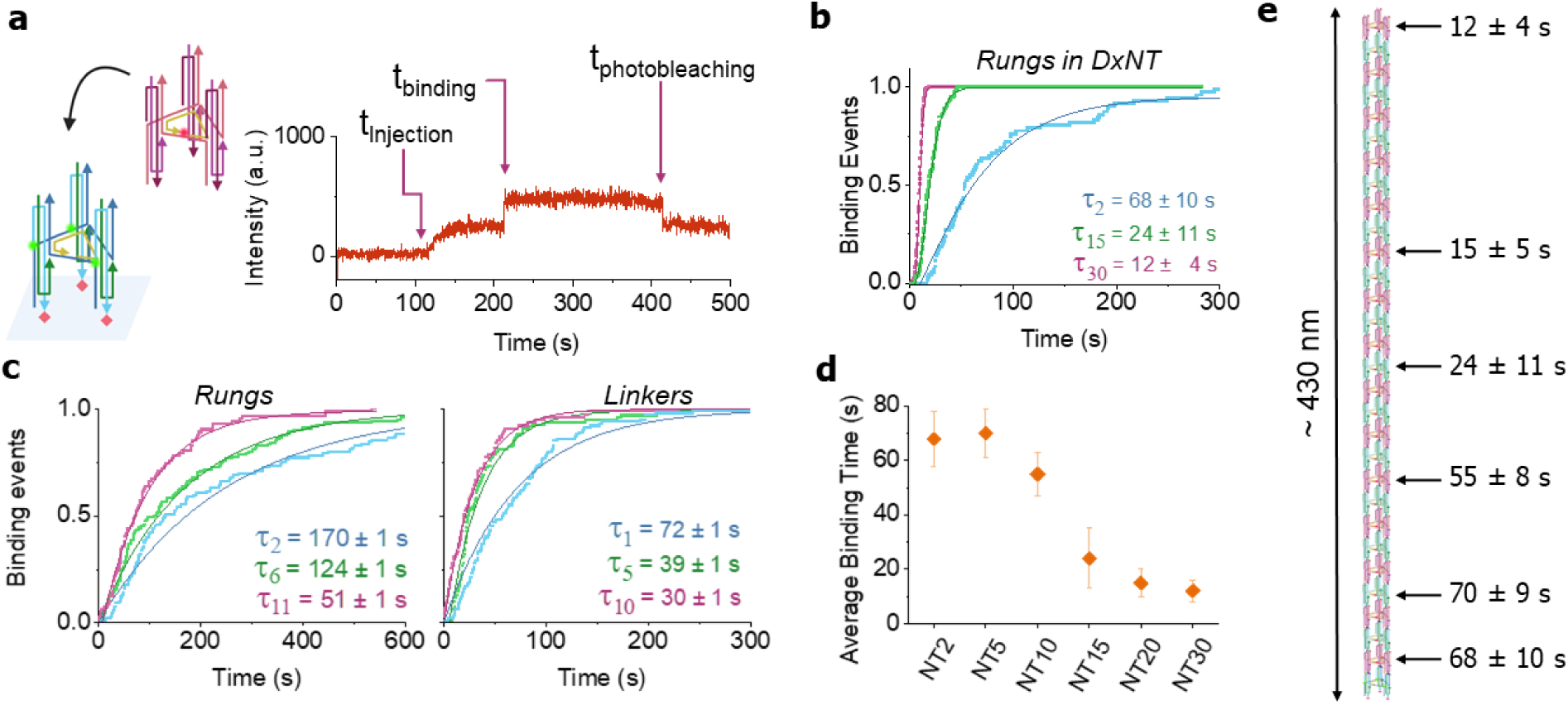
*In situ* binding kinetic experiments. **a.** Cartoon illustrating the arrival of a fluorescently labeled rung (e.g. 2^nd^, 5^th^, 20^th^, etc) onto an otherwise growing nonfluorescent nanotube; the single molecule intensity-time trajectory illustrates an increase in background intensity upon injection gradual rise, followed by a step intensity jump (binding of fluorescently tagged rung) and subsequent photobleaching. **b.** Sample cumulative binding events over time for the addition of Cy5-labelled rungs at the second level (NT2), fifteenth level (NT15), and thirtieth level (NT30) of surface- immobilized DxNTs. The rung concentration was 10 nM in all cases, and the flow rate set at 2 μl./min to minimize structure degradation (see below). Plots were fitted using a gamma cumulative distribution function. **c**. Cumulative binding event plots for rungs (Cy3) and linkers (Cy5) for the wireframe structure by adding the 2^nd^, 6^th^, and 11^th^ units were fitted with a gamma cumulative distribution function to extract the annealing lifetime for each sample. The rung concentration was 10 nM in all cases, and the flow rate set at 3 μl./min to minimize structure degradation (see below). **d.** Average binding time at different heights of the DxNTs. As the separation from the surface increases, annealing time decreases. **e.** The decrease in binding time for DxNT with distance to the surface is illustrated next to the DxNT.

We determined that the binding lifetimes were impacted by proximity to the PEG-coated coverslip surface (Fig. 2d-e, S5-S7). Notably, the addition of units farther away from the surface (*e.g.* at the tenth or thirtieth position) in DxNTs occurred > five-fold faster than additions at the base (*e.g.* at the second position) for these structures, indicating that mass transport down to the coverslip surface in our laminar flow setup is rate limiting. DxNTs have a large persistence length of ∼2.5 µm,^26^ and appeared to stand perpendicular to the surface as revealed by the point spread function of the 25^th^ or 30^th^ labelled rung, appearing out of focus and enlarged compared to surface attached fluorophores; the latter is consistent with structures where the fluorophore distance to the surface is larger than the depth of field, and where the structure pivots around its anchor point. Experimentally, individual rungs were seen to streak across the field of view, under the flowing conditions and when the objective was focused to monitor the 30^th^ labelled rung on the long nanotubes (Fig. S8b). In turn, for experiments monitoring addition at the 2^nd^ level of nascent nanotubes (objective focusing on the surface) individual rungs seen in focus displayed a random walk (Fig. S8a). The slow addition at the base is also consistent with a higher fluid viscosity/PEG meditated restricted accessibility at the interface.^38^

Binding lifetimes recorded for the flexible DNA wireframe structures (50% faster flow rate than above experiments with DxNT) showed the same trend as for DxNT with distance to the surface and additionally illustrated that rung additions were 2- to 3-fold slower than linker additions, following the same trend for the different positions. We attribute these kinetic disparities to the differences in diffusion between the rod-shaped double-stranded linkers (diffusion coefficient, D = 8.6 x 10^-10^ m^2^/s) and the more globular rungs (D = 5.7 x 10^-11^ m^2^/s, details for calculation can be found in the Supporting Information). These differences in diffusion coefficients may partially account for the binding kinetics, as the slower diffusion of the rung is partially mitigated by its trivalent nature. We also note that the overall flexibility of the wireframe design leads to collapsed structures on the surface, limiting the availability of sticky ends for binding.

The kinetic differences we observed along the length of our structures have important implications for surface-bound nanomaterials for sensing and biophysical applications, where mass transport to the surface will significantly influence response time. These factors should be considered when designing stimuli-responsive materials at the solid-liquid interface. Our findings also highlight the usefulness of flow techniques for probing self-assembly dynamics: The control afforded by our automated setup, combined with fine-grained detail provided by single-molecule fluorescence, made it possible to uncover the topographically controlled kinetics in surface-grafted materials that we have observed in this study.

Using the experimentally determined rates of building block addition, we next optimized the duration of reagent delivery, incubation, and washing steps in the automated synthesis. A minimum of 7.5 minutes of flow at rates of 2 μL/min ensured that the 10 μL volume synthesis chamber was replenished with a new solution. Given the lifetimes recorded for DxNT rung addition (≤68 s a), and establishing a minimum hybridization time of 5 lifetimes to ensure yields higher than 99% per hybridization step, we estimated a minimum time of 5.6 minutes for rungs to undergo hybridization (in other words, when reaction times equivalent to 5 lifetimes have elapsed, the remaining unreacted material is found to be (1/e)^5^ ∼ 0.01 or 1%). Importantly, considering that these hybridization reactions are pseudo first order in nature, simply increasing reactant concentrations by one order of magnitude would results in a proportional drop in time of reaction. Using the above reasoning, we optimized the synthesis parameters of DxNTs down to a single cycle of rung A→ buffer → rung B → buffer taking 30 minutes, with the addition of a single level taking approximately 15 minutes with over 96% yield per level (see below for calculation). No additional incubation time was required beyond the time to flow in and wash the reagent, at flow rates of 2 μL/min. Notably, with increasing DxNT length, the five-fold drop observed in annealing lifetimes implies that cycles may be sped up as the structures grow. Care should be taken however to maintain or reduce the flow rates to mitigate structural damage associated with the shear force of the flow (Supplementary Information Section 8.4). Here, all the synthesis steps were achieved automatically using our platform, requiring no additional user time after initial setup.

For flexible DNA wireframe structures, and given the longest lifetimes recorded for linker and rung additions (72 s and 170 s, respectively), we estimated a minimum time of 6 minutes and 15 minutes for linkers and rungs to undergo hybridization. A thorough washing was also incorporated into the synthesis sequence to minimize undesired interactions. Given the structural flexibility and expected lower of reactivity for these structures, ultimately, we opted for longer times for both hybridization steps (a mixture of flow and incubation times to minimize loss of building blocks) as described in the Supplementary Information, Section 9.5. While it is intuitive that longer incubation times would allow for extended hybridization and therefore better yields, the assembly process is conducted at room temperature, allowing for fraying of the base pairs.^39^ This is a major caveat in letting the reaction proceed by incubating for extended times, as degradation, a competing pathway, will occur. Thus, we observed that when nanotubes were left overnight at 4 °C, a fraction could not be extended as revealed by single-molecule photobleaching studies, indicating denaturation (Fig. S14). In our optimized flexible wireframe synthesis protocol, a single cycle of linkers → buffer → rungs → buffer was thus conducted in seventy-five minutes.

### Assembly and in situ characterization

Having established desired synthesis protocols informed by kinetic experiments we next sought to characterize the overall success of the synthesis by quantifying the product yield and its structural integrity utilizing our single molecule/particle imaging capabilities (Fig. 3).

**Figure 3.**
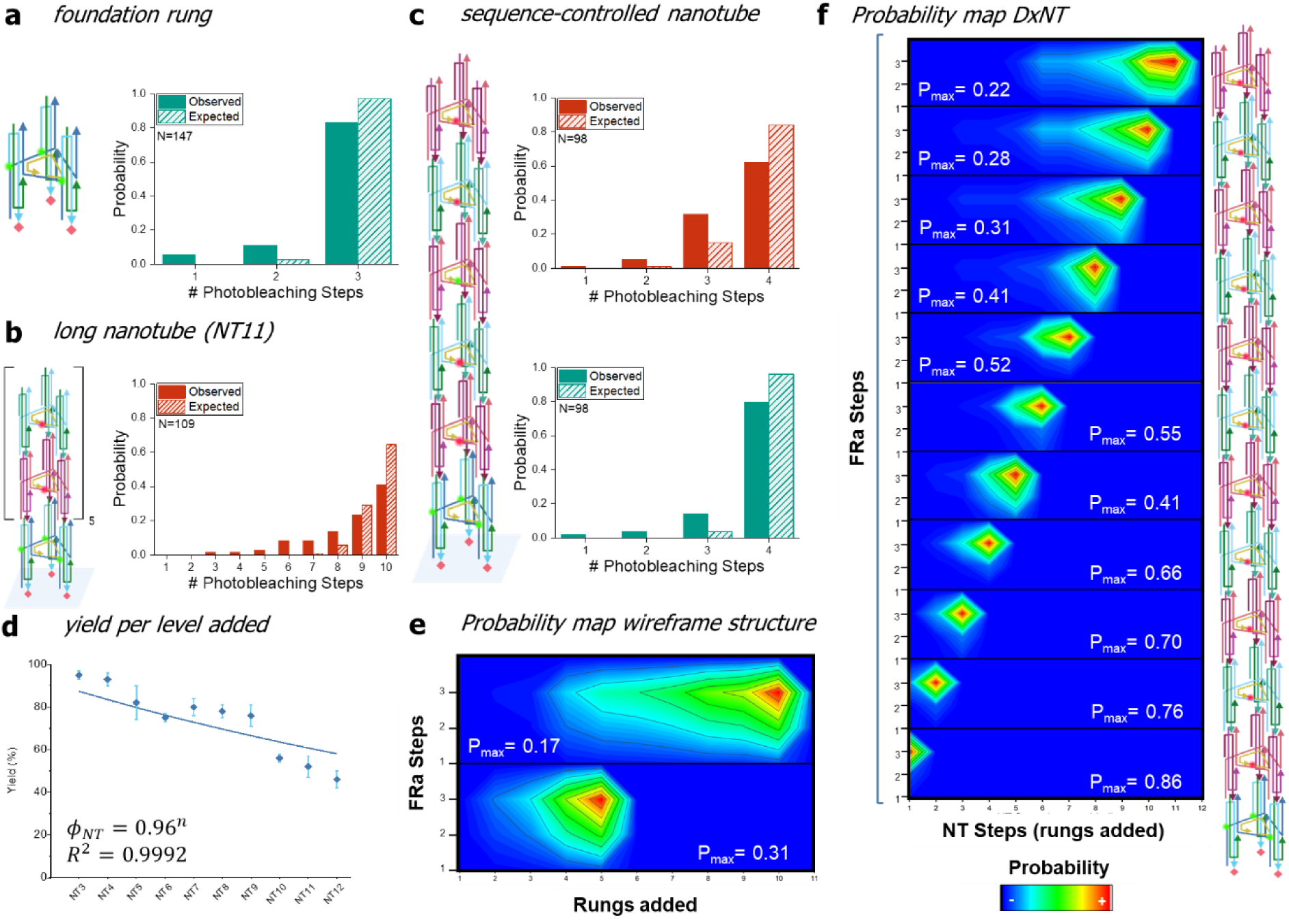
Single-molecule photobleaching experiments demonstrate the ability to form size- and sequence- defined structures. **a.** Photobleaching of the triply-labelled (Cy3) foundation rung and histogram showing the number of photobleaching steps for an entire field of view. The expected values were calculated based on a binominal distribution corresponding to 99.0% labelling efficiency of the Cy3 strand **b.** Structure of an eleven-level (10 expected Cy5-dyes) DxNT and corresponding histogram showing the number of photobleaching steps recorded for the single particles in an entire field of view, with expected values calculated using a binomial distribution corresponding to the 95.7% labelling efficiency of the Cy5-appended strand. **c.** Histograms for dually labelled, sequence-defined nanotube, with a single unique rung at the 4^th^ position. The four green steps represent the triply labelled foundation rung plus the 4^th^ rung, while the four red steps denote four Cy5 labelled rungs. The expected values were calculated using a binomial distribution corresponding to the labelling efficiencies of the two labelled strands – 99.0% for the Cy3-labelled rung and 95.7% for the Cy5-labelled rung. **d.** Calculated yield “ϕ_NT_” for the synthesis of different level “n” DxNTs. A rung addition efficiency “ε”of over 96% as obtained from fitting the resultant data to ϕ_NT_ = ε^n^ . **e.** Two-color correlation for DxNTs formed upon adding n = 5 (5 Cy5 dyes, bottom) and n = 10 (10 Cy5, top) levels to a foundation rung labelled with 3 Cy3 dyes (y-axis). The density map illustrates that the programmed structure is the main product. **f.** Two-color correlation analysis for DxNTs obtained upon adding n = 1 to 11 levels (1 to 11 rungs bearing a single Cy5) to a foundation rung. The desired product bearing 3 Cy3 dyes in the foundation rung (y-axis) and a variable number of Cy5-labelled levels (1-11, x-axis) is shown. The density map indicates that the programmed NT is the main product.

We first ensured the structural fidelity of the foundation rung. These rungs were prepared bearing three fluorophores (either Cy3 or ATTO647N), for structural characterization. Under constant laser illumination in our TIRF microscope setup, each fluorophore bleaches in a stochastic manner, resulting in single particle intensity-time trajectories that appear as a series of steps (analogous to that shown in Fig. 1d for a larger structure), with the number of visible steps indicating the number of incorporated fluorophores. In this way, the formation of the foundation rung and overall NT was assessed at the single-molecule level by leveraging the photobleaching properties of the fluorophores.^35^ The single-molecule photobleaching histogram obtained for the foundation rung agrees with the expected distribution of photobleaching steps based on a binomial distribution. Here, an experimentally determined Cy3 labelling efficiency of 99% (meaning the percentage of strands that actually bear the fluorescent label) was used to compute the expected number of photobleaching steps based on a binomial distribution,^25, 35^ (see Supporting Information for details on labelling efficiencies). A well-formed base was also confirmed via polyacrylamide gel electrophoresis (Fig. S1 and S2). The success of our syntheses (DxNT or flexible wireframe structures, bearing in each case a pre-programmed number of repeat units assembled via the automated method) was next determined – as with the foundation rung - by comparing the observed step count against the theoretical result from the binomial distribution (Fig. S9-11, where labelling efficiencies of 99.0% and 95.7% were experimentally determined for DNA strands used in DxNT rung assembly and labelled with either Cy3 or Cy5, respectively).

To extend our automation to the structural analysis of newly rendered constructs, we developed a linear time-complexity method for detecting intensity steps in single-molecule intensity-time trajectories, based on the binary K-means clustering algorithm,^40^ to quickly screen for synthetic success.^41^ The step detection algorithm operates through iterative binary clustering of the fluorescence versus time trace. We designed an algorithm of linear time complexity, such that the algorithm runtime remains robust to increases in the sampling rate of a photodetector, or other increases to the size of the input data. As shown in Fig. 1f-g, the clustering generated by our algorithm closely matches our manual step counts, with >90 % of traces falling within an error of one count (Fig. 1h), permitting the rapid screening of data sets to estimate synthetic success. Our algorithm is broadly applicable for the rapid assessment of single-molecule photobleaching data to judge synthetic procedures by automatically counting the steps (units) in any given color channel (see Supplementary Information Section 9, and Fig. S15-18 for algorithm details), streamlining the use of single-molecule imaging and analysis to determine synthetic yields. This information can thus be used to e.g. quickly screen different conditions, such as flow rate, to improve assembly procedures in a cyclical workflow.

Qualitatively, an analysis based on counted photobleaching steps, including two-color correlations (Fig. 3e-f), indicates that the nanotubes are formed to the user’s specifications. Comparison of the expected versus experimental probability distributions of photobleaching steps, however, reveals deviations between the theoretical and experimental distributions, where the probability for shorter nanotubes (fewer fluorophores) to occur is larger than anticipated.

To quantify the number of successfully prepared nanotubes, we compared the experimentally obtained probability distribution of dyes in a structure (from photobleaching experiments) and the theoretical expectation based on the binomial model and the experimental labelling efficiency, see Fig. 3. Specifically, we sought to differentiate between 1) complete structures with incomplete labelling, and 2) malformed structures by comparing the various populations within the distribution to the “full count” product. For example, for NT11 in Fig. 3b, the population showing ten steps in the Cy5 channel (the foundation rung bears Cy3 dyes and is thus not probed) must be both correctly assembled and fully labelled, and therefore can be used to assess how many of the nanotubes with nine or fewer steps are fully formed yet lack Cy5 dyes based on the statistics of strand labelling.

The population of malformed DxNTs was estimated by assuming that the experimental distribution of dyes consists of properly formed structures - with their theoretical distribution based on labelling – and truncated structures. By subtracting the theoretical distribution normalized to the yield from the experimental distribution, the truncated components can be retrieved as the difference. Using this method, yields of 85% and 52% were recorded for constructs adding 4 and 10 units, respectively, while a yield of 62% was attained for the sequence-encoded nanotube in Fig. 3c. The correlation of yield for properly formed structures (ϕ_NT_) versus number of steps provided a clear trend wherefrom the efficiency at each cycle (ε) of our synthesis could next be estimated by fitting the experimental results according to ϕ_NT_ = ε^n^, where n denotes the cycle number for the DxNTs. A value of ε = 96% was obtained from fitting the resultant data. Importantly, for flexible wireframe DNA structures it was not possible to estimate the efficiency per cycle as yields (lower compared to DxNT as a function of “n”) did not follow a clear trend with “n”. Here, the putative collapsed morphology and resulting increased steric hindrance may impede hybridization with increasing “n”, i.e. all levels are not equally probable to be formed.

The extent of malformed DxNTs and wireframe structures increased with the flow rate, as revealed by the experimental distribution of number of dyes in our single particle imaging analysis. Slower flow rates (2-3 μL/min or less) resulted in better assembly yields, while faster flow rates (5-15 μL/min) resulted in broader distributions of nanotube length with increased number of smaller structures (Fig. S12-13), suggesting that a parallel degradation pathway in our non-covalent structures is concomitant with nanotube growth, likely due to the mechanical shearing by the fluid. Upon optimizing our procedures through the variation of conditions, we were able to produce nanotubes of varying lengths and sequences (Fig. 3), including asymmetric structures such as our sequence-encoded DNA nanotube (Fig. 3c) where a single unit at the center bearing a Cy3 fluorophore differs from two neighbors above and below (bearing Cy5 in each case).

### DxNT Detachment and surface reutilization

To enable next the selective dislodgement of our newly prepared structures from the surface, we used our automated strategy to insert a unique rung bearing toeholds at the junction with the lower unit. Addition of a fully complementary invading strand elicited detachment of the nanotube from the surface via strand displacement mechanism.^27, 34^ The modified overhang rung (Rb-ov) was added at two different NT lengths (NT2 and NT12) to explore the impact that proximity to the surface may have in dislodgement of the NT at the junction of RB-ov with the lower rung, of interest given the kinetic results of rung annealing (Fig. 2d, see above). Rb-ov was prepared elongating the 3’ sticky end of the “T2-strand’ in each arm with a 10-adenine long sequence overhang (see Fig. 4a).^26^ Addition of the 15 mer complementary invader strand, we posited, would lead to a weakened structure where only three of the initial six bindings would remain between Rb-ov and the lower rung in the DxNT. A DxNT was next prepared adding only Cy3-labelled rungs before the addition of Rb-ov, and subsequently adding the Cy5-labelled Rb-ov followed by additional rung units also bearing Cy5 (Fig. 4b). This arrangement was conceived to monitor in real-time the detachment of red-emitting NTs, leaving behind a green emitting “ nanotube stem”, after the addition of the invader strand.

**Figure 4.**
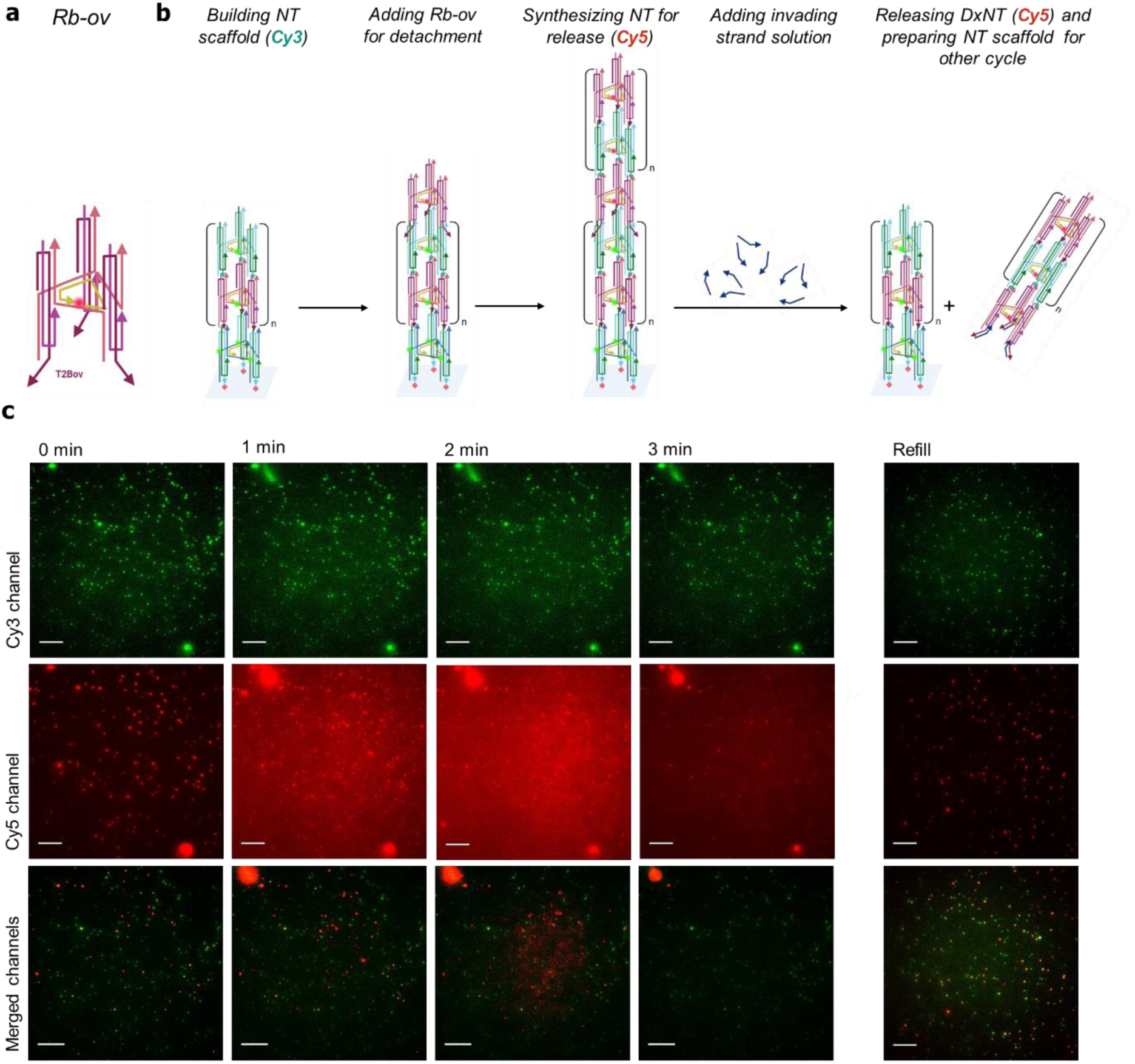
DxNT detachment and surface reutilization. **a**. Design of a modified unit with a 10A overhang at the 3’-end of the T2b strand (**Rb-ov**). **b**. Experimental scheme of detachment experiments. Rb-ov is added to a DxNT at either level 2 or 12 (NT scaffold). NT release happens upon the addition of a completely complementary invading strand. **c.** Example of movie showing DxNT detachment within the first 4 minutes of addition of invading strand (left) and refilling of NT stem after first NT detach experiment (right). Top row shows the Cy3 channel (NT stem), followed by the Cy5 channel (NT to detach), and lastly, bottom row shows the image resulting from merging both channels (where the dynamic range was optimized to assist in visualizing both channels). The disappearance of red signals implies the release of NTs from the surface. After the detachment of DxNTs, an addition of a 10 nM solution of red-labelled rungs can replenish the NT building scaffold. All scale bars= 10 µm.

Imaging studies were conducted sampling the red (departing DxNT segment) and yellow (remaining DxNT segment) emission channels successively (100 ms integration time for each of them). To minimize photobleaching one set of yellow/red frames was acquired per minute during 1 h (60 sets of images). Consistent with controlled detachment of the DxNT at the base of Rb-ov via strand displacement, upon flowing the invading strand (50 μM, 2 μL/min, starting immediately after the first set of images), all particles were observed to disappear from the field of view in the red channel, within three minutes and in a single step (Fig. 4c). In agreement with the second order nature of the reaction, no detachment was observed within 30 min upon flowing 1 μM invading strand. The lack of a stepwise drop in red intensity at the single particle level, and the long survival time (over 20 min) observed for single particles in the red channel in control experiments in the absence of invader strand, imaging under otherwise identical conditions, ruled out photobleaching as the cause of the disappearance of the particles in the red channel. The single particles were still observed in the yellow channel after their disappearance in the red channel, consistent with the lower portion of the DxNT remaining attached to the surface (see Fig. 4b for a 6 min time lapse). Discrete DxNT were seen to detach with the start of the flow, followed by an increased background in the red channel arising from the detached NT in the third-fourth frames as the solution is fully replaced with invading strand. Subsequent frames indicated the red-stained nanotube segments were washed from the field of view as the background decreased to initial values.

To explore the potential for surface reutilization, a fresh solution of Cy5 labelled rung B was next flowed following DxNT detachment. We observed that ∼50% of the remaining DxNT “stems” appearing in the yellow channel bound to the incoming rungs highlighting the potential for the foundation rungs to be utilized over many cycles of DxNT growth and detachment, albeit higher yields of rehybridization are desirable (Fig. 4c).

We note that the detachment of the flexible wireframe DNA structure was not possible either by using a strand displacement mechanism or via separation from streptavidin using a buffer solution containing an excess of biotin. We attribute these failings to steric hindrance and unavailability of the sticky ends due to nanotube collapse.

Importantly, our automated strategy enables the efficient generation of multiple complex structures. Take for example a DxNT construct wherein two different rung units are available, one bearing a yellow emissive dye, the other a red emissive dye. In this case, 2^10^ different nanotube arrangements may be obtained in a 10-rung long nanotube based on the cargo/dye labeling. Here, the total number of possible nanotube arrangements (NT_T_) arises from the ordering of the differing available rungs “r” (*e.g.,* rungs bearing different cargos or labels) along a nascent nanotube. Using probability considerations, one may thus estimate that for a given nanotube structure bearing a pre- defined number of rungs “n” (*e.g.,* 10), NT_T_ increases as a power function (power of ten) on the number of differing rungs NT_T_ = r^n^. Cost, in turn, increases linearly with r. Expanding to a new rung bearing *e.g.,* a green emissive dye, 3^10^ different nanotube arrangements would be obtained, positioning these materials as nanoscale barcodes. By the same token, DNA nanotubes may also be used as biosensors, with the ability to site-specifically add unique building blocks leading to new opportunities for these architectures. Sensing modules such as proteins, aptamers, and antibodies are all compatible with DNA-based self-assembly.

## CONCLUSION

Our custom-made DxNT and flexible wireframe DNA structures are, to our knowledge, the first examples of a fully automated supramolecular synthesis on a solid support. This is a generalizable method for the automated preparation of self-assembled nanostructures with *in situ* single-molecule characterization, allowing for fine-grained detail on the assembly kinetics to be identified upon the variation of self-assembly conditions. This in turn provides an opportunity to eliminate malformed products through the iterative improvement of automated parameters. While we have focused here on the construction of wireframe DNA structures, which are not bound by the size limitations of a scaffold strand, the synthesis and characterization methods described may also find applications in DNA origami and brick-based approaches.^18, 42^ Our microfluidic setup is simple by design and relies on widely available syringe pumps, allowing our methodology to be reproduced in any standard laboratory. Similarly, our screening algorithm can quickly assess synthetic success, making the entire procedure adaptable and efficient. For the rigid DxNT system, the addition of toeholds (DNA overhangs) part way up the nanotube, allowed for the selective dislodgement of our newly prepared structures from the surface upon addition of a complementary invading strand. This strategy will enable the collection of newly prepared nanotubes while also facilitating the recycling of the surface, to upscale production.

The ability to insert unique structural elements into these architectures site-specifically without requiring human intervention or vast numbers of DNA sequences will enable the next generation of surface-grafted biosensing materials with detection at the molecular level. While the scale of production is currently limited, the nanotubes produced herein are suitable for single-molecule assays. We anticipate that these results will provide the foundation for future applications of automated synthesis in supramolecular chemistry and materials science.

## Supporting Information

A detailed materials and methods section is included in the supporting information, which also contains the DNA sequences used, representative single-molecule fluorescence images, additional photobleaching data, a description of our step counting algorithm, and sample traces for all kinetics measurements.

## Acknowledgements

G.C. and H.F.S. are thankful to the National Science and Engineering Research Council of Canada (NSERC), the Canada Foundation for Innovation (CFI), the Fonds de Recherche Nature et Technologies (FRQNT), the New Frontiers in Research Funding (NFRF) and the Canada Institute for Health Research (CIHR) for funding. H.F.S. is thankful to the Canada Research Chairs Program and is a Cottrell Scholar of the Research Corporation. P.I. thanks the Consejo Nacional de Humanidades, Ciencias y Tecnologías (CONAHCyT) for an International Doctoral Scholarship. C.M.P and Y.G. are thankful to NSERC for a CGS-D Scholarship and Vanier Scholarship, respectively. C.M.P. and Y.G. are also thankful to the Drug Discovery and Training Program (CIHR), and Y.G. thanks the NSERC CREATE Bionanomachines Program, for postgraduate scholarships. R.K. is thankful to NSERC for an Undergraduate Summer Research Award. Special thanks to F.J. Rizzuto, D. Saliba, and K.P.F. Janssen for helpful discussions.

## Author contributions

G.C. and H.F.S. conceived the study; C.M.P. and P.I. optimized the setup and performed the single-molecule fluorescence experiments with L.G.; C.M.P and P.I. analyzed the single molecule data with Y.G., G.C. and H.F.S.; P.I. adapted DxNT design for the automated synthesis system, and performed HPLC measurements and analyzed the data to determine the labelling efficiencies. R.K., Y.G., and G.C. designed the step counting algorithm, R.K. wrote the algorithm. Y.G. designed the microfluidic setup and wrote the programs to control the pumps and extract the single-molecule trajectories. G.C. and H.F.S. coordinated and oversaw the project. C.M.P., P.I., G.C. and H.F.S. wrote and edited the manuscript, with input from all the authors.

## Data availability statement

All data supporting the findings of this study are available in the main text and supporting materials. Raw data can be made available upon request.

## Code availability statement

All code can be made available upon request.

## References

1. Caruthers, M. H., The chemical synthesis of DNA/RNA: our gift to science. J. Biol. Chem. 2013, 288 (2), 1420–7.

2. Lutz, J.-F.; Ouchi, M.; Liu, D. R.; Sawamoto, M., Sequence-Controlled Polymers. Science 2013, 341 (6146), 1238149.

3. Calin, O.; Eller, S.; Seeberger, P. H., Automated polysaccharide synthesis: assembly of a 30mer mannoside. Angew. Chem. Int. Ed. 2013, 52 (22), 5862–5865.

4. Joseph, A. A.; Pardo-Vargas, A.; Seeberger, P. H., Total synthesis of polysaccharides by automated glycan assembly. J. Am. Chem. Soc. 2020, 142 (19), 8561–8564.

5. Hoogenboom, R.; Meier, M. A.; Schubert, U. S., Combinatorial methods, automated synthesis and high-throughput screening in polymer research: past and present. Macromol. Rapid Commun. 2003, 24 (1), 15–32.

6. Pan, X.; Lathwal, S.; Mack, S.; Yan, J.; Das, S. R.; Matyjaszewski, K., Automated synthesis of well-defined polymers and biohybrids by atom transfer radical polymerization using a DNA synthesizer. Angew. Chem. Int. Ed. 2017, 56 (10), 2740–2743.

7. Li, J.; Ballmer Steven, G.; Gillis Eric, P.; Fujii, S.; Schmidt Michael, J.; Palazzolo Andrea, M. E.; Lehmann Jonathan, W.; Morehouse Greg, F.; Burke Martin, D., Synthesis of many different types of organic small molecules using one automated process. Science 2015, 347 (6227), 1221–1226.

8. Blair, D. J.; Chitti, S.; Trobe, M.; Kostyra, D. M.; Haley, H. M. S.; Hansen, R. L.; Ballmer, S. G.; Woods, T. J.; Wang, W.; Mubayi, V.; Schmidt, M. J.; Pipal, R. W.; Morehouse, G. F.; Palazzolo Ray, A. M. E.; Gray, D. L.; Gill, A. L.; Burke, M. D., Automated iterative Csp3– C bond formation. Nature 2022, 604 (7904), 92–97.

9. Buitrago Santanilla, A.; Regalado Erik, L.; Pereira, T.; Shevlin, M.; Bateman, K.; Campeau, L.-C.; Schneeweis, J.; Berritt, S.; Shi, Z.-C.; Nantermet, P.; Liu, Y.; Helmy, R.; Welch Christopher, J.; Vachal, P.; Davies Ian, W.; Cernak, T.; Dreher Spencer, D., Nanomole-scale high-throughput chemistry for the synthesis of complex molecules. Science 2015, 347 (6217), 49–53.

10. Gesmundo, N. J.; Sauvagnat, B.; Curran, P. J.; Richards, M. P.; Andrews, C. L.; Dandliker, P. J.; Cernak, T., Nanoscale synthesis and affinity ranking. Nature 2018, 557 (7704), 228–232.

11. de Greef, T. F. A.; Meijer, E. W., Supramolecular polymers. Nature 2008, 453 (7192), 171–173.

12. Brunsveld, L.; Folmer, B. J. B.; Meijer, E. W.; Sijbesma, R. P., Supramolecular Polymers. Chem. Rev. 2001, 101 (12), 4071–4098.

13. Lo, P. K.; Karam, P.; Aldaye, F. A.; McLaughlin, C. K.; Hamblin, G. D.; Cosa, G.; Sleiman, H. F., Loading and selective release of cargo in DNA nanotubes with longitudinal variation. Nat. Chem. 2010, 2 (4), 319–328.

14. Lehn, J.-M., From supramolecular chemistry towards constitutional dynamic chemistry and adaptive chemistry. Chem. Soc. Rev. 2007, 36 (2), 151–160.

15. Seeman, N. C.; Sleiman, H. F., DNA nanotechnology. Nat. Rev. Mater. 2017, 3, 17068.

16. Yan, H.; LaBean, T. H.; Feng, L.; Reif, J. H., Directed nucleation assembly of DNA tile complexes for barcode-patterned lattices. Proc. Natl. Acad. Sci. USA 2003, 100 (14), 8103–8108.

17. Yan, H.; Feng, L.; LaBean, T. H.; Reif, J. H., Parallel Molecular Computations of Pairwise Exclusive-Or (XOR) Using DNA “String Tile” Self-Assembly. J. Am. Chem. Soc. 2003, 125 (47), 14246–14247.

18. Rothemund, P. W., Folding DNA to create nanoscale shapes and patterns. Nature 2006, 440 (7082), 297–302.

19. Fu, J.; Yan, H., Controlled drug release by a nanorobot. Nat. Biotechnol. 2012, 30 (5), 407–408.

20. Acuna, G. P.; Möller, F. M.; Holzmeister, P.; Beater, S.; Lalkens, B.; Tinnefeld, P., Fluorescence Enhancement at Docking Sites of DNA-Directed Self-Assembled Nanoantennas. Science 2012, 338 (6106), 506–510.

21. Li, J.; Johnson-Buck, A.; Yang, Y. R.; Shih, W. M.; Yan, H.; Walter, N. G., Exploring the speed limit of toehold exchange with a cartwheeling DNA acrobat. Nature Nanotechnology 2018, 13 (8), 723–729.

22. Chidchob, P.; Sleiman, H. F., Recent advances in DNA nanotechnology. Curr. Opin. Chem. Biol. 2018, 46, 63–70.

23. Veneziano, R.; Ratanalert, S.; Zhang, K.; Zhang, F.; Yan, H.; Chiu, W.; Bathe, M., Designer nanoscale DNA assemblies programmed from the top down. Science 2016, 352 (6293), 1534.

24. Sanderson, K., Bioengineering: What to make with DNA origami. Nature 2010, 464 (7286), 158–9.

25. Hariri, A. A.; Hamblin, G. D.; Gidi, Y.; Sleiman, H. F.; Cosa, G., Stepwise growth of surface-grafted DNA nanotubes visualized at the single-molecule level. Nat. Chem. 2015, 7 (4), 295–300.

26. Luo, X.; Saliba, D.; Yang, T.; Gentile, S.; Mori, K.; Islas, P.; Das, T.; Bagheri, N.; Porchetta, A.; Guarne, A.; Cosa, G.; Sleiman, H. F., Minimalist Design of Wireframe DNA Nanotubes: Tunable Geometry, Size, Chirality, and Dynamics. Angew. Chem. Int. Ed. 2023, 62 (44), e202309869.

27. Platnich, C. M.; Hariri, A. A.; Rahbani, J. F.; Gordon, J. B.; Sleiman, H. F.; Cosa, G., Kinetics of Strand Displacement and Hybridization on Wireframe DNA Nanostructures: Dissecting the Roles of Size, Morphology, and Rigidity. ACS Nano 2018, 12 (12), 12836–12846.

28. Gidi, Y.; Bayram, S.; Ablenas, C. J.; Blum, A. S.; Cosa, G., Efficient One-Step PEG-Silane Passivation of Glass Surfaces for Single-Molecule Fluorescence Studies. ACS Appl. Mater. Interfaces 2018, 10 (46), 39505–39511.

29. Platnich, C. M.; Rizzuto, F. J.; Cosa, G.; Sleiman, H. F., Single-molecule methods in structural DNA nanotechnology. Chem. Soc. Rev. 2020, 49 (13), 4220–4233.

30. Wang, W.; Yu, S.; Huang, S.; Bi, S.; Han, H.; Zhang, J.-R.; Lu, Y.; Zhu, J.-J., Bioapplications of DNA nanotechnology at the solid–liquid interface. Chem. Soc. Rev. 2019, 48 (18), 4892–4920.

31. Chen, Z.; Lichtor, P. A.; Berliner, A. P.; Chen, J. C.; Liu, D. R., Evolution of sequence-defined highly functionalized nucleic acid polymers. Nat. Chem. 2018, 10 (4), 420–427.

32. Ulbrich, M. H.; Isacoff, E. Y., Subunit counting in membrane-bound proteins. Nat. Methods 2007, 4 (4), 319–321.

33. Casanova, D.; Giaume, D.; Moreau, M.; Martin, J.-L.; Gacoin, T.; Boilot, J.-P.; Alexandrou, A., Counting the number of proteins coupled to single nanoparticles. J. Am. Chem. Soc. 2007, 129 (42), 12592–12593.

34. Simmel, F. C.; Yurke, B.; Singh, H. R., Principles and Applications of Nucleic Acid Strand Displacement Reactions. Chem. Rev. 2019, 119 (10), 6326–6369.

35. Hariri, A. A.; Hamblin, G. D.; Hardwick, J. S.; Godin, R.; Desjardins, J.-F.; Wiseman, P. W.; Sleiman, H. F.; Cosa, G., Stoichiometry and Dispersity of DNA Nanostructures Using Photobleaching Pair-Correlation Analysis. Bioconj. Chem. 2017, 28 (9), 2340–2349.

36. Cordes, T.; Vogelsang, J.; Tinnefeld, P., On the Mechanism of Trolox as Antiblinking and Antibleaching Reagent. J. Am. Chem. Soc. 2009, 131 (14), 5018–5019.

37. Glembockyte, V.; Cosa, G., Redox-Based Photostabilizing Agents in Fluorescence Imaging: The Hidden Role of Intersystem Crossing in Geminate Radical Ion Pairs. J. Am. Chem. Soc. 2017, 139 (37), 13227–13233.

38. Andrian, T.; Pujals, S.; Albertazzi, L., Quantifying the effect of PEG architecture on nanoparticle ligand availability using DNA-PAINT. Nanoscale Adv. 2021, 3 (24), 6876–6881.

39. Andreatta, D.; Sen, S.; Pérez Lustres, J. L.; Kovalenko, S. A.; Ernsting, N. P.; Murphy, C. J.; Coleman, R. S.; Berg, M. A., Ultrafast Dynamics in DNA: “Fraying” at the End of the Helix. J. Am. Chem. Soc. 2006, 128 (21), 6885–6892.

40. Forgy, E. W., Cluster analysis of multivariate data: efficiency versus interpretability of classifications. Biometrics 1965, 21, 768–769.

41. Shapiro, S. S.; Wilk, M. B., An analysis of variance test for normality (complete samples). Biometrika 1965, 52 (3/4), 591–611.

42. Hong, F.; Zhang, F.; Liu, Y.; Yan, H., DNA origami: scaffolds for creating higher order structures. Chem. Rev. 2017, 117 (20), 12584–12640.

