## Supplemental Material for "Automated Synthesis of Wireframe DNA Nanotubes"

### Table of contents:


1. Materials
2. Oligonucleotides
    - 2.1. Wireframe Structure
    - 2.2. DxNT
3. Preparation of DNA nanotube subunits
    - 3.1. Wireframe structure subunits
    - 3.2. DxNT subunits
    - 3.3. Gel electrophoresis of foundation rung for wireframe structure
    - 3.4. Gel electrophoresis of rungs for DxNT
4. Automated synthesis setup
    - 4.1. Materials and components summary
    - 4.2. Cable connections and computer control of the pumps
    - 4.3. Standard cycles
5. Single-molecule surface preparation
6. Single-molecule TIRF
    - 6.1. Image and data analysis
7. Single-molecule binding kinetics
    - 7.1. Time selection
    - 7.2. Binding curves
    - 7.3. Iterative cycle improvement
    - 7.4. Stokes-Einstein calculations for rung and linker
    - 7.5. Speed calculations
8. Single-molecule photobleaching
    - 8.1. Labelling efficiency
    - 8.2. Photobleaching steps histograms
    - 8.3. Binding errors
    - 8.4. Flow rate impact on synthesis yield
    - 8.5. Structure integrity over time
9. Step-count screening algorithm
    - 9.1. Simulating intensity-time profiles
10. References

### *1. Materials*

Acetic acid, boric acid, EDTA, urea, magnesium chloride, GelRed, tris(hydroxymethyl)aminomethane (Tris), D(+) glucose, β-mercaptoethanol, and streptavidin were purchased from Aldrich or BioBasic Inc. Nucleoside (1000 Å)-derivatized LCAA CPG solid supports with loading densities of 25–40 μmol/g, Sephadex G-25 (super fine DNA grade), and reagents for automated DNA synthesis were used as purchased from BioAutomation. Acrylamide (40%)/bis-acrylamide 19:1 solution was purchased from BioShop. For TIRF-M surface sample preparation, poly(ethylene glycol) silanol MW 5000 (mPEG-Sil) and biotin-PEG-Sil were purchased from Laysan Bio, Inc. Imaging chamber components were custom-made, purchased from Grace Bio-Labs. TEAA buffer is composed of 50 mM triethylamine and 50 mM Acetic Acid with a pH of 8.0. TBE buffer is composed of 90 mM Tris and boric acid and 1.1 mM EDTA with a pH of ~8.3. TAMg buffer is composed of 45 mM Tris and 12.5 mM $MgCl_2$ with a pH of ~7.8 adjusted by glacial acetic acid.

### *2. Oligonucleotides*

DNA strands were ordered from Integrated DNA Technologies (IDT). Labelled strands were bought HPLC purified. For unlabeled strands, desalted products were purified by running them on 12% polyacrylamide/8M urea polyacrylamide gels (20 x 20 cm vertical Hoefer 600 electrophoresis unit) at a constant current of 30 mA for 1-2 hours using 1 x TBE as the running buffer. Following electrophoresis, the gels were illuminated with a UV lamp (254 nm) and the bands were excised. The resulting gel pieces were then crushed and incubated in 10 mL of sterile water at 55 °C for 16 hours. Samples were then dried to 1.5 mL, desalted using size exclusion chromatography (Sephadex G-25) and quantified by UV/visible spectroscopy with a NanoDrop One$^{C}$ Spectrophotometer and using IDT's extinction coefficient at 260 nm.

***2.1. Wireframe structure.*** All sequences are given in the 5' to 3' direction.

| **Strand name** | **Length** | **Sequence** |
|---|---|---|
| **V** | 113 | CTCAGCAGCGAAAAACCGCTTTACCACATTCGAGGCACGTTGTACGT CCACACTTGGAACCTCATCGCACATCCGCCTGCCACGCTCTTAGCATA GGACGGCGGCGTTAAATA |
| **C1** | 84 | CGGTGCATTTCGACGGTACTTCGTACAACGTGCCTCGAATGTAGAGC GTGGCAGGCGGATGTGAAGCAGTTGCAGCGTACTCGT |
| **C2** | 63 | TCGGCAGACTAATACACCTGTCGATGAGGTTCCAAGTGTGGATAGCT AGGTAACGGATTGAGC |
| **R1F** | 54 | TGGCGACGGTCGCGAGGTTATTTAACGCCGCCGTCCTATGCTTTGTAA AGCGGT |
| **R2F** | 50 | TGGCGACGGTCGCGAGGTGCTCAATCCGTTACCTAGCTCCAGTACCGT CG |
| **R3F** | 54 | TGGCGACGGTCGCGAGGTACGAGTACGCTGCAACTGCTACCAGGTGT ATT |
| **Cy5Biotin** | 18 | Cy5-ACCTCGCGACCGTCGCCA-Biotin |
| **Cy3Biotin** | 18 | Cy3-ACCTCGCGACCGTCGCCA-Biotin |
| **ATTO647NBiotin** | 18 | ATTO647N-ACCTCGCGACCGTCGCCA-Biotin |
| **Cy3R1** | 22 | TGCAACTGCTACCAGGTGTATT-Cy3 |
| **ATTO647NR1** | 22 | TGCAACTGCTACCAGGTGTATT-ATTO647N |
| **R2** | 22 | TTACCTAGCTCCAGTACCGTCG |
| **R3** | 22 | GTCCTATGCTTTGTAAAGCGGT |
| **LS1** | 62 | TTTTCGCTGCTGAGGTAAGCCTTCGGCGAGCATCTATCTATGTCTCCG TATTTAAC GCCGCC |
| **LS1*** | 34 | CGGAGACATAGATAGATGCTCGCCGAAGGCTTAC |

| | | |
|---|---|---|
| **LS1* ATTO647N** | 44 | ATTO647N-TTTTTTTTTTCGGAGACATAGATAGATG CTCGCCGAAGGCTTAC |
| **LS2** | 62 | AGTCTGCCGACACAGAGATCAGTCGGAAGCATAATATCTTATGTTCG TGATAACG AGTACGC |
| **LS3** | 62 | AAATGCACCGCACAGAGATCAGTCGGAAGCATAATATCTTATGTTCG TGATAGCT CAATCCG |
| **LS2/3*** | 42 | TATCACGAACATAAGATATTATGCTTCC GACTGATCTCTGTG |

***2.2. DxNT structure.*** All sequences are given in the 5' to 3' direction.

| Strand name | Length | Sequence |
|---|---|---|
| **C3** | 63 | TCAACTGCTCATCCTATATGGTCAACTGCTCA TCCTATATGGTCAACTGCTCATCCTATATGG |
| **C3 Cy3** | 63 | Cy3-TCAACTGCTCATCCTATATGGTCAACTGCTCA TCCTATATGGTCAACTGCTCATCCTATATGG |
| **C3 Cy5** | 63 | Cy5-TCAACTGCTCATCCTATATGGTCAACTGCTCA TCCTATATGGTCAACTGCTCATCCTATATGG |
| **HA** | 70 | GTTTCCACTATAGAAGAGCGTAATCTGAGCAGTTGACCATATAGGAC GAAGTATGAATACCAGATGTCGT |
| **T1A** | 60 | ACTGATATCTTGGTCCATTTAGGATTACGCTCTTCTATAGTGCATTATC TATGTAAGTAT |
| **T1A Biotin** | 60 | Biotin-ACTGATATCTTGGTCCATTTAGGATTACGCTCTTC TATAGTGCATTATCTATGTAAGTAT |
| **T2A** | 59 | CTAAATGGACCAAGATAATCTGGTATTCATACTTCGATACTTACATAG ATAATGTTGCT |
| **T2A Cy3** | 59 | Cy3-CTAAATGGACCAAGATAATCTGGTATTCATACTTCG ATACTTACATAGATAATGTTGCT |
| **HB** | 70 | TCAGTCACTATAGAAGAGCGTAATCTGAGCAGTTGACCATATAGGAC GAAGTATGAATACCAGATAGCAA |
| **T1B** | 60 | GAAACTATCTTGGTCCATTTAGGATTACGCTCTTCTATAGTGCATTAT CTATGTAAGTAT |
| **T2B** | 59 | CTAAATGGACCAAGATAATCTGGTATTCATACTTCGATACTTACATAG ATAATGACGAC |
| **T2B-10A** | 69 | CTAAATGGACCAAGATAATCTGGTATTCATACTTCGATACTTACATAG ATAATGACGACAAAAAAAAAA |
| **SD-10T** | 15 | TTTTTTTTTTGTCGT |

#### *3. Preparation of DNA nanotube subunits*

#### *3.1. Wireframe structure subunits*

To prepare the foundation rungs, the constituent nine strands were combined in equimolar amounts in 1 x TAMg buffer. This mixture was then annealed at 95 °C then slowly cooled down to 4 °C over 6 h to give the foundation rung in quantitative yield.

Subsequent rungs were formed by combining the six constituent strands in equimolar amounts in 1 x TAMg buffer. This mixture was annealed at 95 °C then slowly cooled down to 4 °C over 6 h to give the rung in quantitative yield.

Linking strands were formed by combining the two constituent strands in equimolar amounts in 1 x TAMg buffer. This mixture was annealed at 95 °C then slowly cooled down to 4 °C over 6 h to give the linking strand in quantitative yield.

#### *3.2. DxNT subunits*

There is a minimum of three building blocks necessary to perform the solid phase synthesis of the DxNT. Rungs are formed by annealing four different strands. The foundation rung (FRa) is comprised of HA, T1A Biotin, T2A Cy3, and C3. Rungs A and B are formed by their respective H, T1, T2 strands and C3 Cy5. For the triangular cross-section NT, three equivalents of the H, T1, and T2 strands are needed per equivalent of the C3 strand. All mixtures were annealed at 95 °C then slowly cooled down to 4 °C over 14 h to give the linking strand in quantitative yield.

***3.3. Gel electrophoresis of foundation rung for wireframe structure:*** Linking strands were formed by combining the two constituent strands in equimolar amounts in 1 x TAMg buffer. This mixture was annealed at 95 °C then slowly cooled down to 4 °C over 6 h to give the linking strand in quantitative yield.

. While the linking strands and rungs reported here have previously been characterized by our groups, the foundation rung is a new structure. Notably, the foundation rung used here is composed purely of unmodified DNA and requires no further purification.

Assembly of the foundation rung was verified using native polyacrylamide gel electrophoresis (PAGE). Gels were run with 15% glycerin in 1 x TAMg buffer at 4 °C at a constant voltage of 100 V and stained with GelRed in 1 x TAMg prior to imaging.

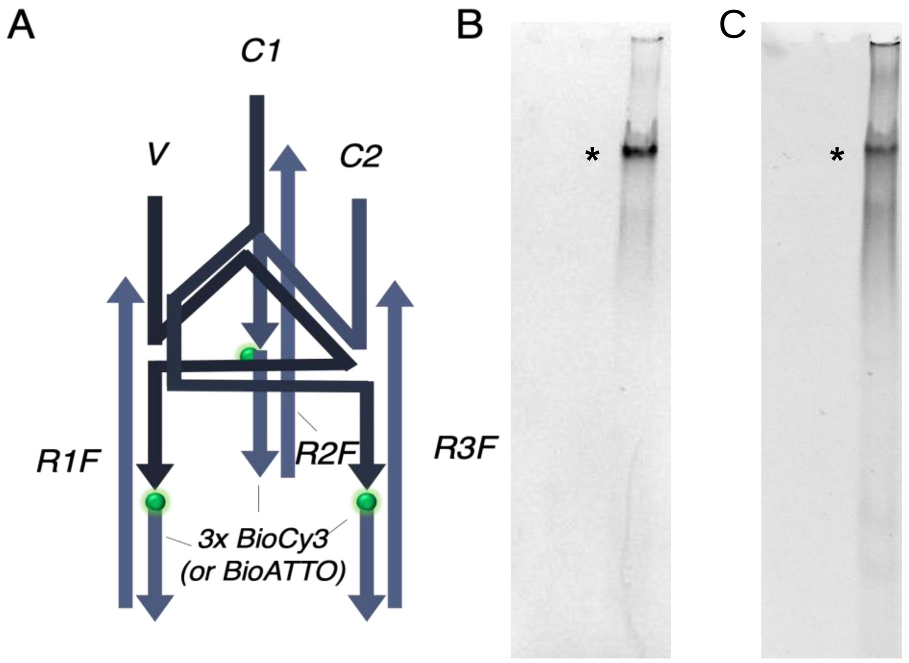


**Figure S1. Assembly of the foundation rung.** A 6% native PAGE in 1xTAMg is shown to illustrate the assembly of the foundation rung, with the major product being the fully formed structure. The sample was imaged in both the (B) Cy3 channel and the (C) GelRed channel.

***3.4. Gel electrophoresis of rungs for DxNT.*** The purity of individual strands was established by denaturing PAGE. This gel ran with 4 M urea in 1x TBE buffer at room temperature at a constant voltage of 100 V and stained with GelRed in 1 x TBE prior to imaging.

Assembly of the rungs was verified using native PAGE. Gels were run with 15% glycerin in 1 x TAMg buffer at 4 °C at a constant voltage of 100 V and stained with GelRed in 1 x TAMg prior to imaging.

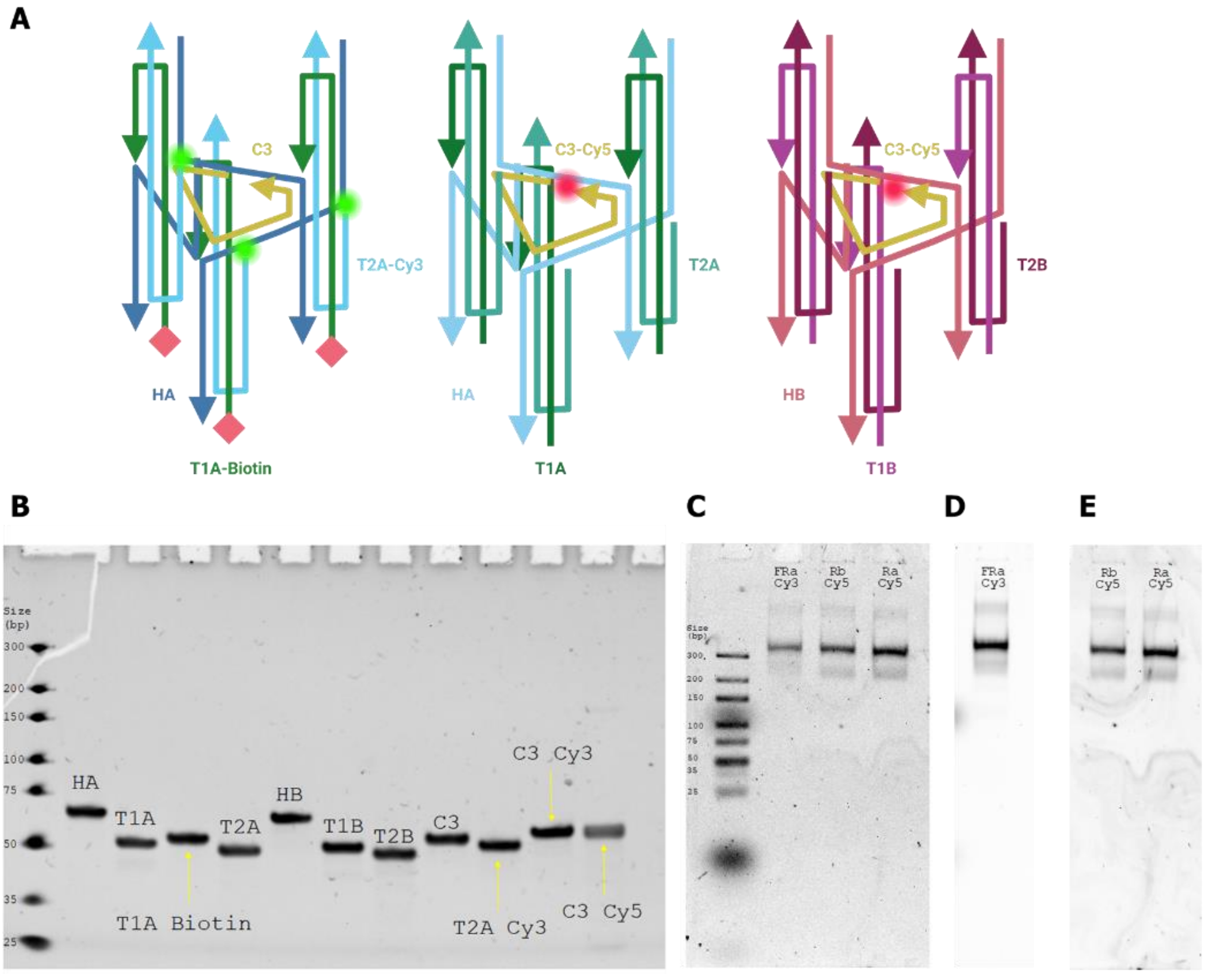


**Figure S2. Assembly of each rung for DxNT.** (**A**) Rung design with all sequences for foundation rung, rung A, and rung B. (**B**) 12% denaturing PAGE for all strands imaged after GelRed staining to verify DNA strands are clean for assembly. (**C**) 6% native PAGE in 1xTAMg is shown to illustrate the assembly of the foundation rung, rung A, and rung B after annealing. For all, the major product is the fully formed structure. The sample was imaged in the GelRed channel, (**D**) Cy3 channel, and the (**E**) Cy5 channel.

#### *4. Automated synthesis setup*

##### *4.1. Materials and components summary*

The automated synthesis setup consisted of several computer-controlled syringe pumps allowing the control of flow rates and duration of pump action. 1 mL Syringes were loaded with buffer solutions containing buffer only, linkers, and rungs, for the wireframe structure. Buffer, Ra, and Rb solutions for DxNT. Syringes were connected with FEP tubbing. The tubing-syringe fitting assembly consisted of a ferrule, a Nut, and a female adapter for each syringe. All tubing exiting the multiple syringes converged to a 4-port cross assembly. The exit port of the manifold was directed to the coverslip access ports. The exit port of the coverslip was directly connected to FEP tubing and directed to a waste container.

| COMPONENT | NUMBER | COST PER UNIT (USD) | TOTAL COST | SOURCE | MATERIAL |
|---|---|---|---|---|---|
| **COMPUTER-CONTROLLED SYRINGE PUMPS (PHD 2000, HARVARD APPARATUS)** | 3 | 475.00** | 1,425.00 | ebay.com | n.a. |
| **1 mL SYRINGES (NORM-JECT, AIR-TITE) (100 PACK)** | 1 | 12.80* | 12.80* | https://ca.vwr.com/store/product?keyword=53548-001 | Polypropylene barrel and a Polyethylene plunger |
| **FEP TUBING (OD = 1/32”, ID = 0.1 mm, VICI JOUR)** | 3 | 20.00 | 60.00 | https://www.vicijour.com/products/fep-tubing | Fluorinated Ethylene Propylene |
| **FERRULE (1/4-28, 1/32” OD, P-248X, IDEX) (10 PACK)** | 1 | 44.00* | 44.00* | https://scipro.com/product/idex-p-248x-fittings-super-flangeless-ferrule-m6-with-ss-ring-1-32-od-10pk/ | n.a. |
| **NUT (1/4-28 FLAT BOTTOM, P-252X, IDEX) (10 PACK)** | 1 | 19.30* | 19.30* | https://www.coleparmer.com/i/idex-super-flangeless-nut-standard-knurl-gray-pom-1-32-or-1-16-od-tubing-1-4-28-flat-bottom-10-pk/0202162 | n.a. |
| **FEMALE SYRINGE ADAPTER (P-628, IDEX)** | 3 | 6.80* | 20.40* | https://scipro.com/product/idex-p-628-connector-female-luer-adapter-to-female-union/ | n.a. |
| **4-PORT CROSS ASSEMBLY (ETFE, P-634, IDEX)** | 1 | 50.00* | 50.00* | https://www.coleparmer.com/p/idex-low-pressure-crosses/72887?Ntt=IDEX+P-634 | n.a. |
| **PRESS FIT TUBING CONNECTORS (20 PACK)** | 1 | 29.00 | 29.00 | https://gracebio.com/product/press-fit-tubing-connectors-460003/ | n.a. |
| **GRACE BIO-LABS HYBRIWELL CHAMBER (0.010 PC + 1LSA); 1-13mm X 4mm ID, 18mm X 18 mm OD (100 PACK)** | 1 | 237.00 | 237.00 | Custom chamber, direct quote from https://gracebio.com/ | Polycarbonate |
| **SQUARED GLASS COVERSLIPS (25 X 25)mm** | 1 | 15.70* | 15.70* | https://www.fishersci.ca/shop/products/fisherbrand-cover-glasses-squares-8/12542c | Borosilicate Glass |

*Prices in CAD exchanged to USD.
**Refurbished; the model used is currently discontinued.

***4.2. Cable connections and computer control of the pumps.*** Pumps were controlled by a computer using serial RS-232/USB communication. A RJ11 “straight Thru” cable was connected to the exit port of the pump and converted into a USB port using two adapters (RJ11/female DB9) and (male DB9/USB). Syringe pumps (PHD 2000, Harvard Apparatus) were independently connected to the computer and controlled using a code written in Python. Communication was achieved using the freely available “Pumpy” package (https://github.com/tomwphillips/pumpy). A custom Python-based script was written to control flow sequences and the number of cycles and can be made available upon request.

***4.3. Standard cycle.***

**Wireframe Structure.**

**Lines flush.** Each line corresponding to (1) Buffer, (2) Rungs, and (3) Linkers was initially filled with the corresponding solution at a 5 μL/min rate for 10 minutes.

**Cycles.** The cycle initiates with the addition of linkers to the foundation rung (3 μL/min rate for 10 minutes, 5-minute incubation) followed by the addition of buffer (3 μL/min rate for 20 minutes). Next, the rung solution is added (3 μL/min rate for 10 minutes, 10-minute incubation) to the previously hybridized linkers. The cycle ends with the addition of buffer solution (3 μL/min rate for 20 minutes). Each cycle takes approximately 95 minutes. The number of cycles is added as an input in the Pumps-controlling software and can be changed as per convenience. Changes to the cycle order as well as the addition of new pumps to deliver new structures is a straightforward process. The concentration was 20 nM for rung and linker additions.

**DxNT.**

**Lines flush.** Each line corresponding to (1) Buffer, (2) Rb, and (3) Ra was initially filled with the corresponding solution at a 5 μL/min rate for 10 minutes.

**Cycles.** The cycle initiates with the addition of Rb to the foundation rung (3 μL/min rate for 7 minutes, 2-minute incubation) followed by the addition of buffer (3 μL/min rate for 7 minutes). Next, the Ra solution is added (3 μL/min rate for 7 minutes, 2-minute incubation) to the previously hybridized Rb. The cycle ends with the addition of buffer solution (3 μL/min rate for 7 minutes). Each cycle takes approximately 32 minutes, and it adds 2 levels. The number of cycles is added as an input in the Pumps-controlling software. The concentration was 10 nM for rung solutions.

***5. Single-molecule surface preparation***

Surface preparation was performed by following the procedure reported by Gidi *et al.*[1] Glass coverslips were soaked in piranha solution (30% $H_2O_2$ solution and concentrated $H_2SO_4$, in a 1:3 v/v ratio) and sonicated for one hour. Coverslips were then etched by sonication for 30 minutes in a 0.5 M aqueous solution of NaOH (temperature maintained between 20 °C and 30 °C). Etched coverslips were rinsed three times with water (Hyclone molecular biology grade) then sonicated in water for fifteen minutes. This procedure was then repeated with acetone (HPLC grade) to remove any surface-bound water molecules. Surface passivation with poly(ethylene glycol) silane was then carried out one coverslip at a time. The coverslip was dried under a nitrogen stream then a silicon mold was placed over the coverslip. A 99:1 (w/w) mixture of poly(ethylene glycol) silane (MW 5000) and biotin-poly(ethylene glycol) silane (MW 5000) was then dissolved in dry DMSO (25% PEG-silane w/w). The coverslip was preheated in an oven at 90 °C for 5 min then the freshly prepared PEG-silane/DMSO solution was added. After 15 minutes in the oven, the silicon mold was removed, and the coverslip washed thoroughly with Hyclone water and dried under a stream of nitrogen.

Imaging chambers (~ 10 μL) were formed by pressing a Grace Bio-Labs polycarbonate film with an adhesive gasket onto the PEG-silane-coated coverslips. Two silicone connectors were glued onto the pre-drilled holes of the film to serve as inlet and outlet ports. Prior to imaging, 10 μL of a 1 mg/mL streptavidin solution was added to the surface and left to incubate for 10 min. The excess streptavidin was then washed with 50 μL 1 x TAMg buffer in Hyclone water prior to adding the sample to the surface.

#### *6. Single-molecule TIRF*

Single molecule fluorescence imaging was carried out on a total internal reflection fluorescence (TIRF) microscopy setup consisting of an inverted microscope (Nikon Eclipse Ti) equipped with a motorized illuminator unit and Ti-ND6 Perfect Focus System (PFS). Samples were excited with the evanescent wave of either a 561 nm or 647 nm diode laser output of a monolithic laser combiner (Agilent Technologies, MLC-400B). The beam position was adjusted to attain total internal reflection through an oil-immersion objective (Nikon CFI SR Apochromat TIRF 100x, NA = 1.49). In all cases, the laser beam was passed through a multiband clean-up filter (ZET405/488/561/647x, Chroma Technology) and coupled into the microscope objective using a multiband beam splitter (ZT405/488/561/647rpc, Chroma Technology). Fluorescence light was spectrally filtered with an emission filter (ZET405/488/561/647m, Chroma Technology). For imaging of Cy3, an additional emission filter (ET600/50m, Chroma Technology) was used. During kinetics experiments, an additional emission filter for red dyes (Cy5 and ATTO647N) was used (ET705/72m, Chroma Technology).

The camera was controlled using Micro-Manager software (Micro-Manager 1.4.13, San Francisco, CA), capturing 16-bit 512 x 512 pixel images with an exposure time of 100 ms. All movies were recorded onto a 512 x 512 pixel region of a back-illuminated electron-multiplying charge-coupled device (EMCCD) camera (iXon X3 DU-897-CS0-#BV, Andor Technology). The microscope was controlled using the software NIS element from Nikon.

##### *6.1. Image and data analysis*

We observed typically ~150-200 spots corresponding to single structures, over one field of view (~$82^2$ $\mu m^2$). We utilized a Matlab-based algorithm to perform single-molecule data analysis. Single-molecule detection was based on the IDL algorithm utilized by the Ha group.[2] Diffraction-limited spots (DLSs) are mapped via threshold analysis, and the clustered spots are rejected. Further analysis was performed with an in house written algorithm. Intensity-time single-molecule trajectories are extracted as follows: Initially, the positions of the DLSs were mapped. Then, Intensity-time single-molecule trajectories are extracted. Background correction was performed for each fluorescence spot at every frame, considering its surrounding background.

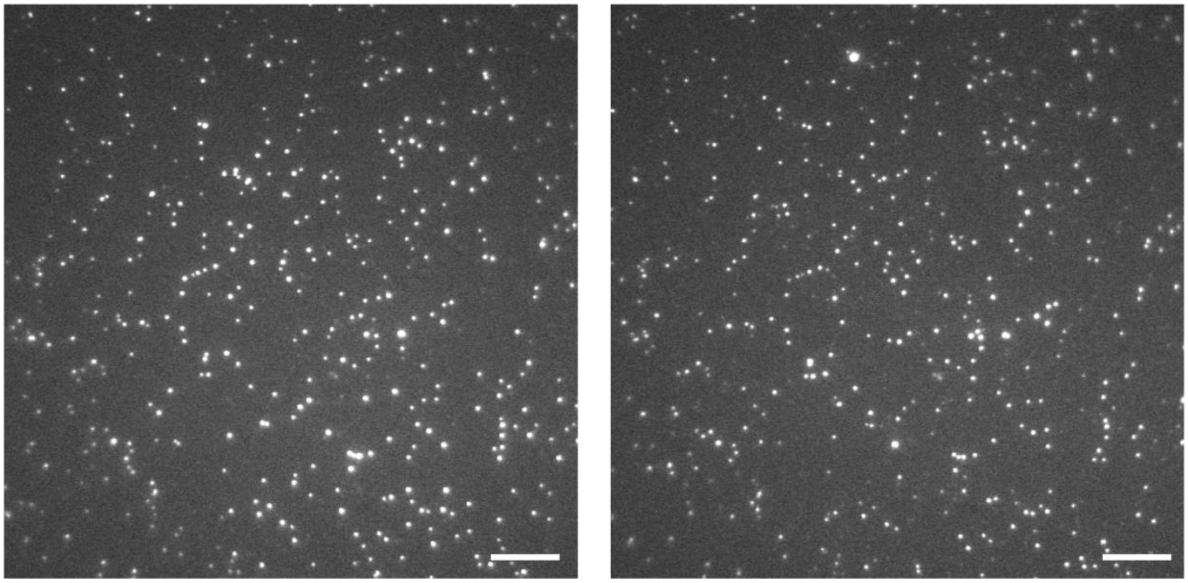


**Figure S3. Representative single molecule images of newly assembled DNA nanostructures.** TIRF images taken directly after automated synthesis cycles of red (ATTO647N) foundation rungs with ten green (Cy3) rungs assembled on top. The low background of the images indicates that the buffer rinsing steps have been adequate to remove all freely floating constructs in solution. The good correlation between green and red images indicates that binding is specific to the structures and there are negligible non-specifically surface bound fluorophores following washing cycles. The foundation rung was inserted at a concentration of 100 pM. Subsequent rungs and linkers were added at a concentration of 20 nM. Addition and washing cycles were described in section 5.2. Imaging conditions: 100 ms acquisition, 7 mW 647 nm laser, 6 mW 561 nm laser. Scale bars = 10 μm.

#### *7. Single-molecule binding kinetic experiments*

In order to reduce photobleaching events, an oxygen scavenger solution was prepared consisting of a triplet quencher (2 mM Trolox from 200 mM freshly prepared stock in HPLC-grade MeOH) and an oxygen scavenger system (D(+)glucose 3% w/v, glucose oxidase 0.1 mg/mL, and catalase 0.02 mg/mL) in 1 x TAMg buffer. This solution was injected onto the coverslip surface and allowed to incubate for at least 15 minutes prior to image acquisition.[4, 5]

During acquisition, a solution of either fluorescently labelled rungs or linkers was prepared (20 or 10 nM in fresh oxygen scavenger solution). Either of these solutions was then flowed through the imaging chamber at a flow rate of 3 μL/min (mimicking the optimized synthesis conditions) using a syringe pump (Harvard Instruments).

***7.1. Time Selection.*** For binding kinetic experiments, time count started when the background increased due to flow of labeled subunits. Binding time is considered when there is a single step intensity hike.

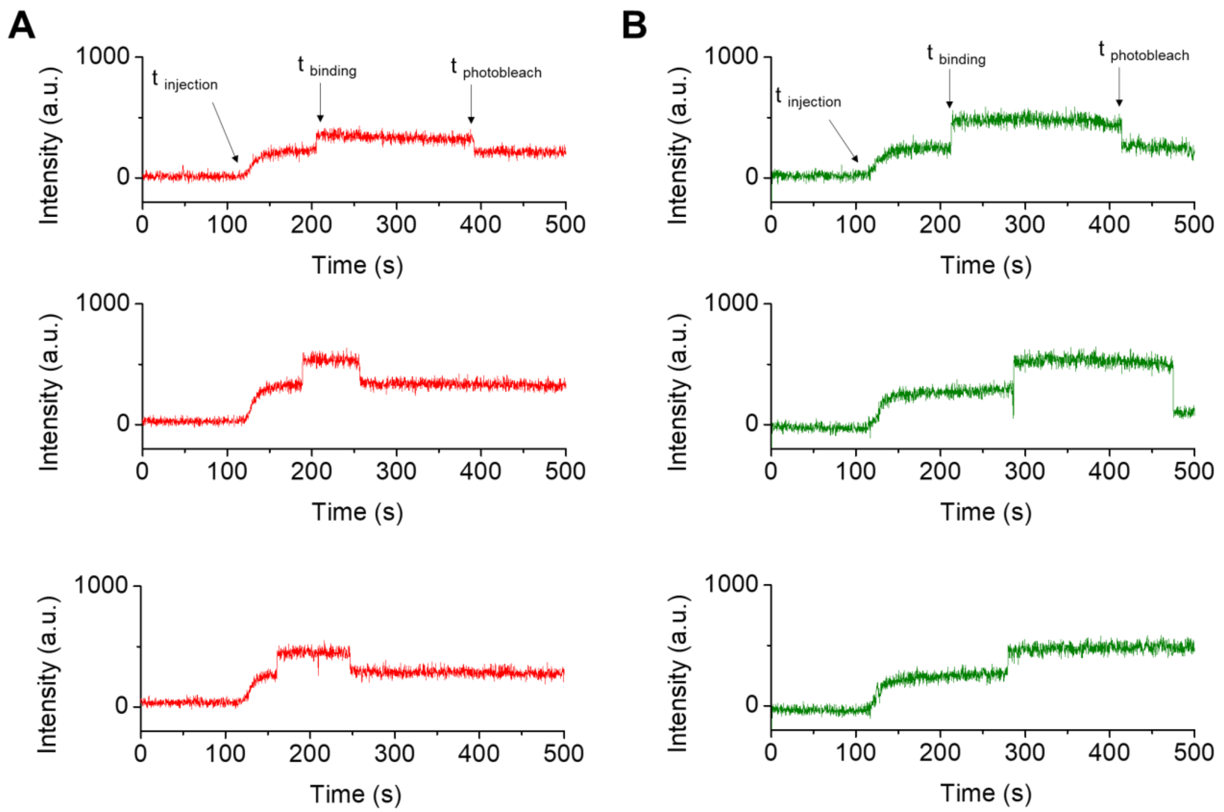


**Figure S4. Non-background corrected single-molecule trajectories.** Trajectories are shown for the addition of the first set of (**A**) linkers, labelled with ATTO647N and (**B**) rungs, labelled with Cy3. When the labelled DNA structures flow into the chamber, their fluorescence causes an increase in the background, highlighted in yellow, after which hybridization occurs, resulting in a single step increase in intensity.

#### *7.2. Binding Kinetic Curves.*

**Wireframe structure.** Plots of cumulative binding events as a function of time were fitted to a gamma cumulative distribution function calculating the average binding time of the subunit added (rungs, linkers).

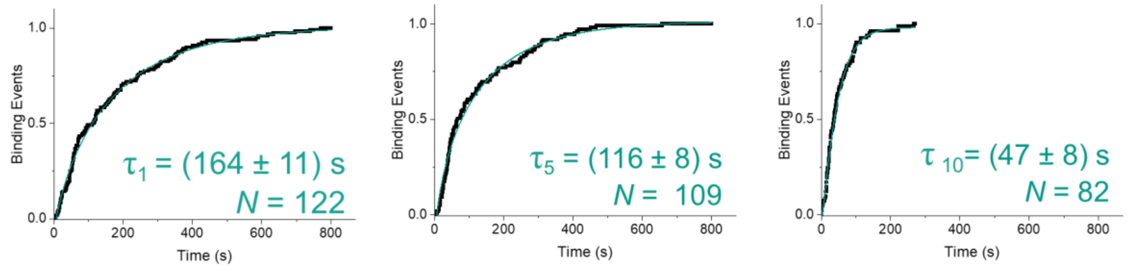


**Figure S5. Kinetics of addition for rungs.** Cumulative binding events as a function of time, fitted to a gamma cumulative distribution function to extract the lifetime for the binding of (**A**) 1st rung, (**B**) 5th rung, and (**C**) 10th rung.

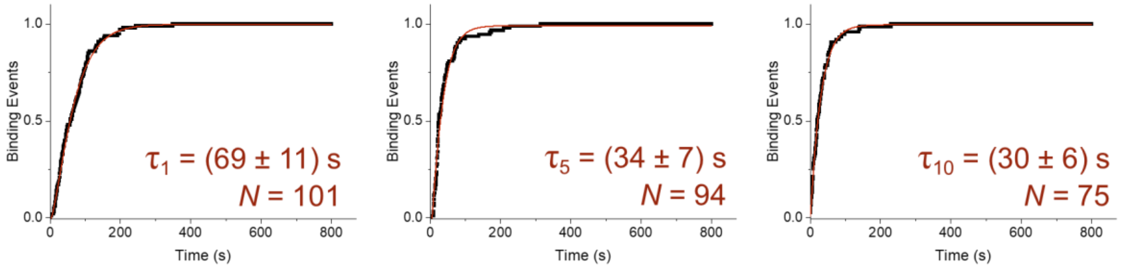


**Figure S6. Kinetics of addition for linkers.** Cumulative binding events as a function of time, fitted to a gamma cumulative distribution function to extract the lifetime for the binding of (**A**) 1st set, (**B**) 5th set, and (**C**) 10th set of linkers.

**DxNT.** Plots of cumulative binding events as a function of time were fitted to a gamma cumulative distribution function calculating the average binding time of the subunit added. Binding times clearly decreased as the height of the NT grows away from the surface.

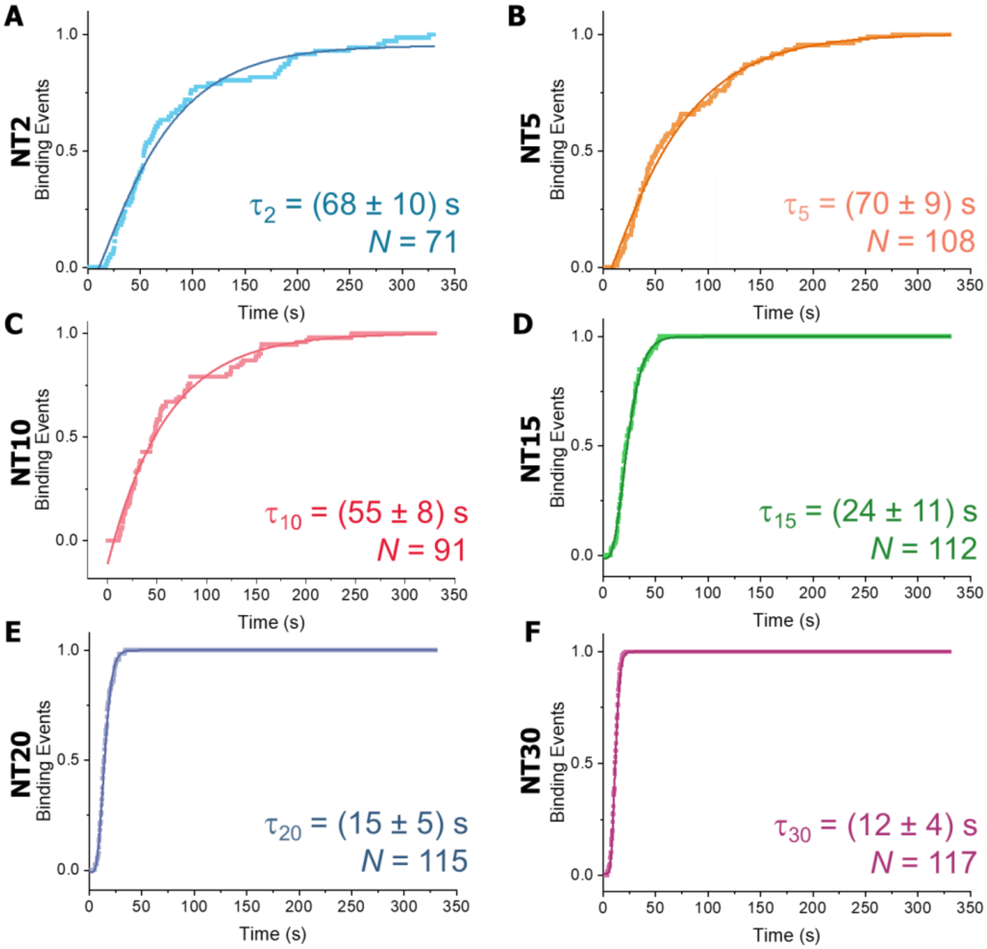


**Figure S7. Kinetics of addition for the n-th rung.** Cumulative binding events as a function of time, fitted to an gamma cumulative distribution function to extract the lifetime of rung addition for the (**A**) 2nd level, (**B**) 5th level, (**C**) 10th level, (**D**) 15th level, (**E**) 20th level, and (**F**) 30th level. Panel **G** displays the average binding time in comparison to NT length.

*7.3. Stokes-Einstein calculations for rung and linker*

To rationalize the kinetic differences between the rung and the linking strand we used the Stokes-Einstein equation to approximate the diffusion coefficients of these two components. The rung was approximated as a sphere with a radius ($R_0$) of 3.74 nm (based on the geometry of the rung itself). The linker was approximated as oblate (rod-shaped),[8] with the major axis ('a') being the length of the double-stranded unit (~ 21 nm) and the minor axis ('b') as the width of the double helix (~ 2 nm). While this calculation does not consider the hydrodynamic radius of the components, it still enables a comparison between the rungs and the linkers.

In the case of the DxNT, all units can be considered roughly the same size and shape, where the major axis of the oblate can be considered as the length of the rung (~14.3 nm) and the minor axis as the width of the NT (~6 nm).

$$D\ (sphere) = \frac{k_B T}{6\pi\mu R_0}$$

$$D\ (oblate) = \frac{k_B T}{6\pi\mu\left(\frac{\sqrt{a^2-b^2}}{tan^{-1}\left[\sqrt{\frac{a^2-b^2}{b^2}}\right]}\right)}$$

where $\mu$ is the viscosity of water at room temperature (1.00 x $10^{-3}$ Pa•s).

*7.4. Speed Calculations*

As stated in section 7.2, the average time to bind the last building block decreases as it grows away from the surface, or said in another way, by increasing its height. Due to the laminar flow within the chamber, mass transport is dependant on the height of the construct. In the case of the DxNT, NT height went from ~28.6 nm for NT2, all the way to ~430 nm for NT30.

Looking at the behaviour of labelled rungs around the NT before attaching, it is very clear that units closest to the surface move according to a random walk, since they are the least affect by the flow. However, the flow is evident for taller structures that move in a straight line lining up to the flow.

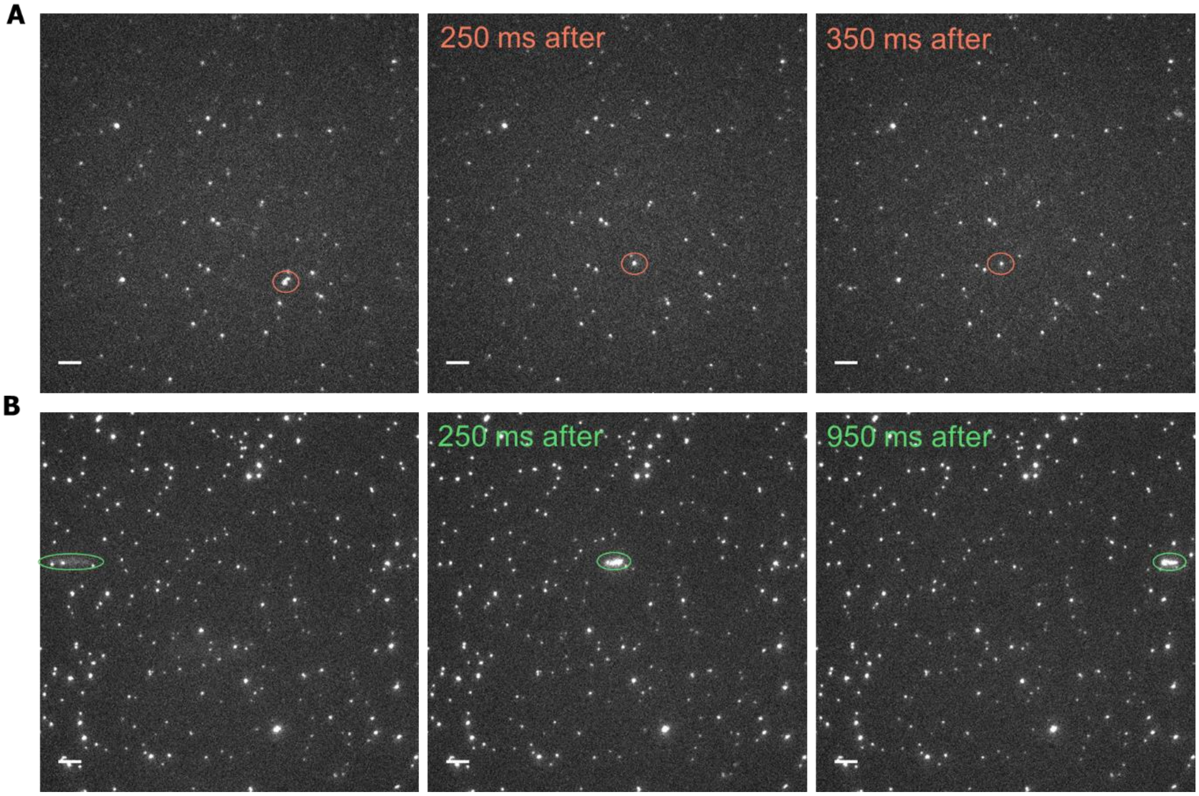


**Figure S8. Movement of binding rungs at the surface and above.** (**A**) Three frames (50 ms each) showing no particular directional movement for run B binding to foundation run to yield NT2. (**B**) Three frames (50 ms each) showing directional movement of rungs binding at the NT30 level. Widefield fluorescent movies were taken while flowing oxygen scavenger solution at 100 µL/min. 5 µm scale bars.

#### *9. Single-molecule photobleaching*

To reduce blinking events associated to *e.g.* fluorophore intersystem crossing, a solution of 2 mM Trolox in 1 x TAMg was injected into the coverslip prior to photobleaching.[3-5] While recording the photobleaching process over time for each bound foundation rung or nanotube, fluorescence images of an ~ $82^2$ $\mu m^2$ area were acquired on an EMCCD camera (100 ms per frame).

**Wireframe structure**. A total of 1200 frames (120 s acquisition time) were typically collected for the ATTO647N emission (foundation rung photobleaching, 647 nm laser excitation, 6-7 mW power output measured from the objective) whereas 2000 frames (200 s acquisition time) were usually collected for Cy3 emission (full nanotube, 561 nm laser excitation, 6-8 mW power output measured from the objective).

**DxNT**. A total of 8000 frames (800 s acquisition time) were typically collected for the Cy5 emission (full nanotube, 647 nm laser excitation, 5 mW power output measured from the objective) whereas 4000 frames (400 s acquisition time) were usually collected for Cy3 emission (foundation rung photobleaching, 561 nm laser excitation, 3 mW power output measured from the objective).

#### *9.1. Labelling efficiency measurements*

The labelling efficiency of each fluorophore-appended DNA strand was measured using reverse phase high-performance liquid chromatography using a Hamilton PRP-1 column (150 mm x 4.1 mm, 10 µm) on an Agilent Infinity 1260 instrument. Each experiment was conducted using a gradient from 3 to 70% of ACN over 30 minutes. By comparing the area of the peak corresponding to the correctly labelled strands against the sum of all peaks with a signal at 260 nm (DNA), the labelling efficiency is obtained.

$$\%\text{Labelling} = 100\% * \frac{A_{labelled\ strand}}{A_{labelled\ strand} + A_{unlabelled\ strand}}$$

***9.2. Photobleaching steps histograms.*** For all structures, yield was calculated by normalizing the expected probability of the product to the calculated perfectly labelled product (100%) and subtracting the malformed nanotubes' probability from it.

On the next figures, the white bars thus represent nanotubes which are fully formed, but incompletely labelled, whereas the solid bars (green or red) denote structures which are truncated or broken. The population of truncated nanotubes increases for longer nanotube lengths, which we attribute to the shear force generated by the laminar flow farther from the coverslip surface.

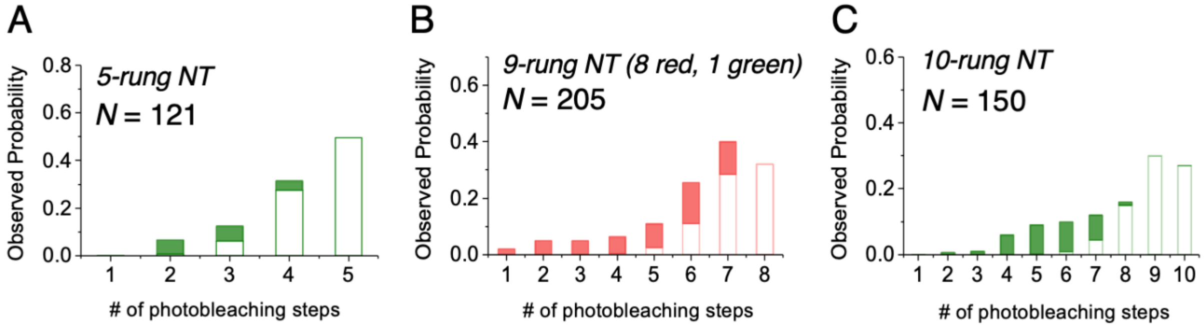


**Figure S9. Properly formed versus truncated constructs for wireframe structure.** Structures were assembled with (**A**) 5 rungs (**B**) 9 rungs (of which 8 were red, and one green, not shown here) and (**C**) 10 cycles. Based on a binomial distribution assuming 90% labelling efficiency, a population of nanotubes that have fewer-than-expected steps is inevitable. Here, we sought to separate the unlabelled population from the malformed by normalizing the distributions with respect to the number we observed of the "full count" product: For the five-rung nanotube in (**A**), for example, those showing five steps must be both correctly assembled and fully labelled.

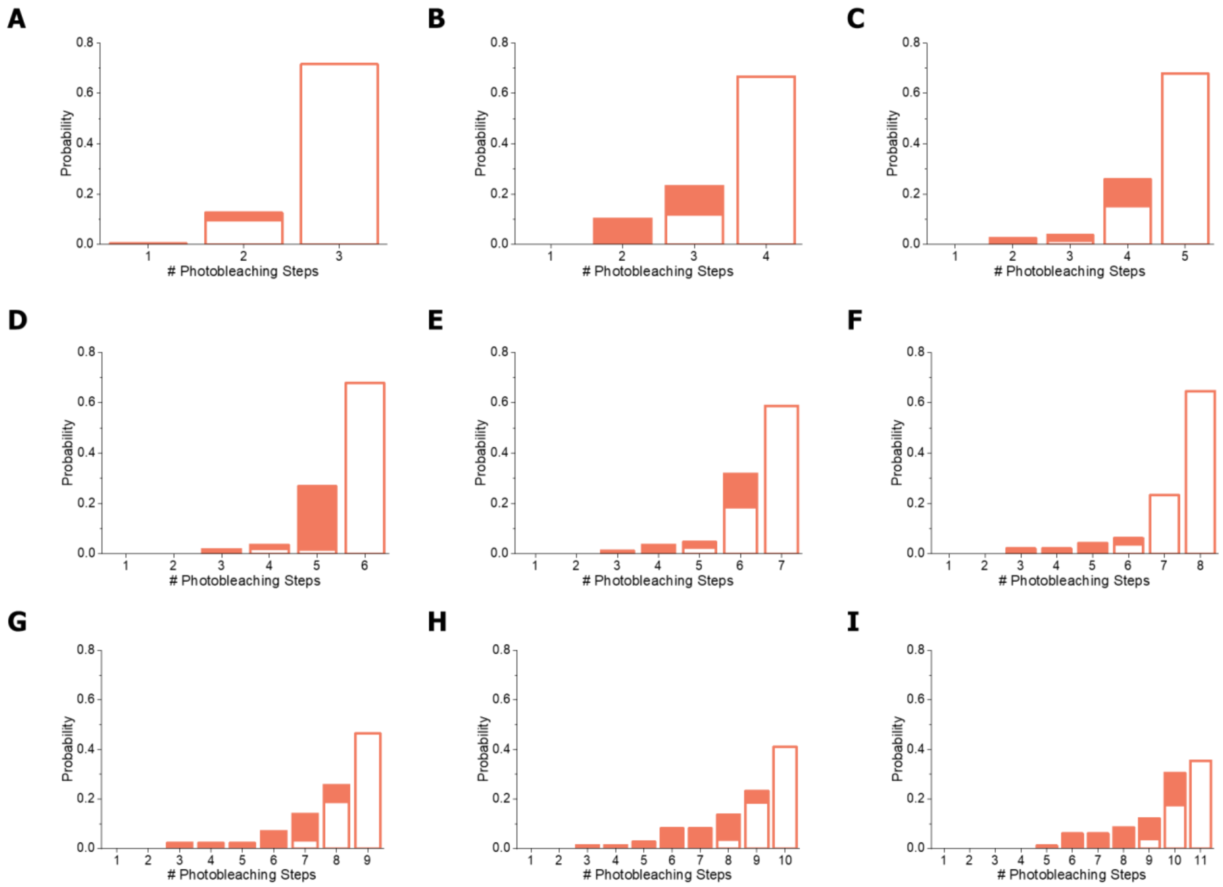


**Figure S10. Properly formed versus truncated nanotubes for DxNT.** Structures were assembled until the (**A**) 4$^{th}$ level, (**B**) 5$^{th}$ level, (**C**) 6$^{th}$ level, (**D**) 7$^{th}$ level, (**E**) 8$^{th}$ level, (**F**) 9$^{th}$ level, (**G**) 10$^{th}$ level, (**H**) 11$^{th}$ level, and (**I**) 12$^{th}$ level. Based on a binomial distribution assuming 95.7% labelling efficiency.

***9.3. Binding errors.*** For the wireframe structure, the probability contour maps show populations different from the main product. These can be attributed to truncated structures and labelling errors.

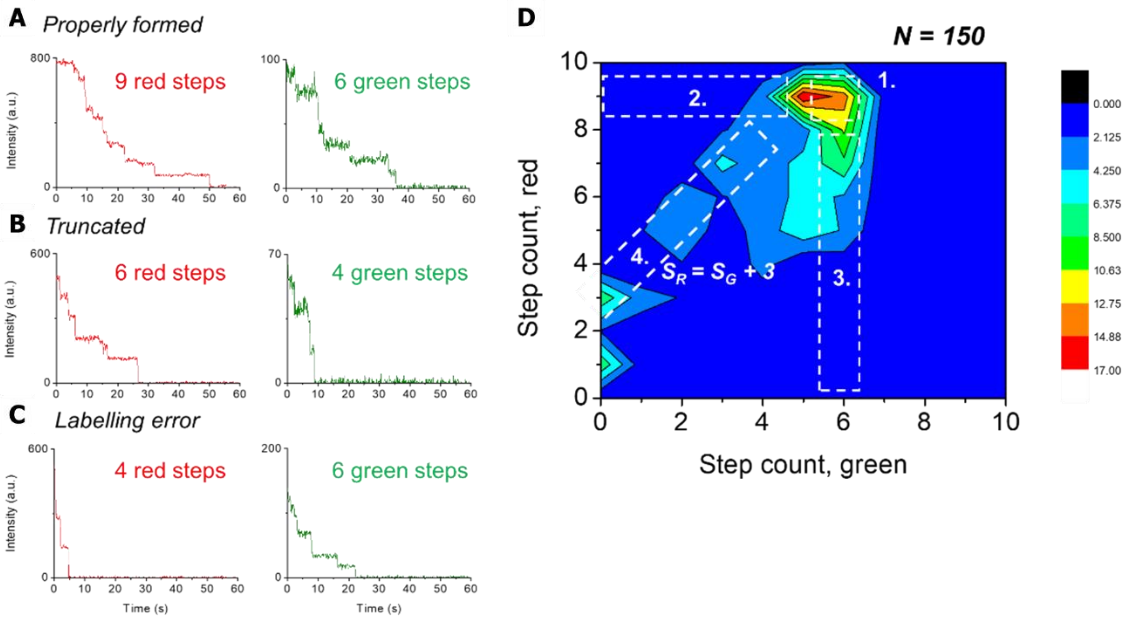


**Figure S11. Two-color correlation analysis for well-formed nanotubes with ATTO647N-labelled foundation rung, six Cy3-labelled rungs, and six ATTO647N-labelled linkers.** (**A**) Cartoon of the structure. Properly formed nanotubes show 9 steps in the red channel and 6 in the green channel (**B**), corresponding to highlighted zone E1 on panel E). Truncated nanotubes (**C**) result from synthesis failures, will have fewer steps in both channels, falling along line E4 in panel **E**. Labelling errors (**D**) result in a mismatch in the number of red and green photobleaching steps and fall along lines E2 and E3 in panel **E**. (**E**) Correlation plot of the number of steps on the green vs the red channel for each individual nanotube in a population of 150 constructs.

***9.4. Flow rate impact on synthesis yield.*** In order to optimize every part of the synthesis protocol, different flow rates were tested. However, at higher flow rates, nanotube breakage occurs, shearing off some of the rungs and resulting in higher populations of smaller nanotubes.

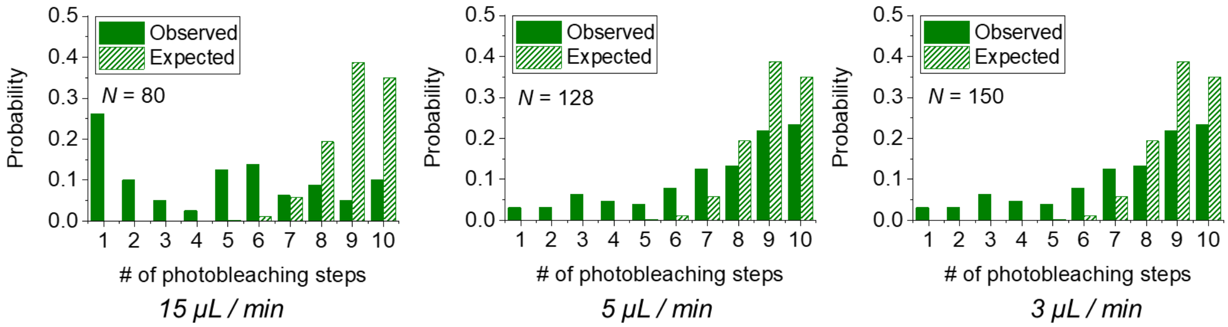


**Figure S12. Effect of flow rate on wireframe structure.** Single-molecule photobleaching histograms for 10-rung structures, with each rung bearing a Cy3 dye, for differing flow rates. As we decrease the flow rate, the population with < 4 rungs drops off significantly, demonstrating a reduction of breakage.

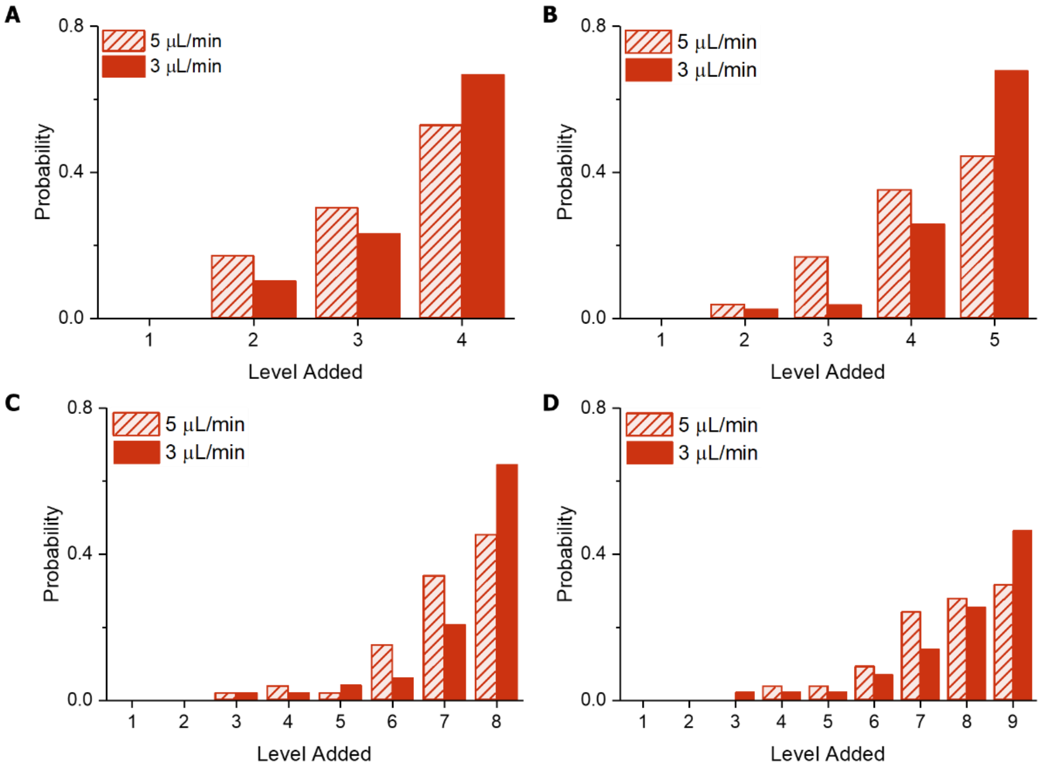


**Figure S13. Effect of flow rate on DxNT structure.** Single-molecule photobleaching histograms for (**A**) NT5, (**B**) NT6, (**C**) NT9, (**D**) NT10, flowing each level at 3 (full color) and 5 μL/min (striped). At 5 μL/min there was already significant breakage, thus no higher injection rates were tried.

***9.5. Construct integrity overnight.*** For the wireframe structure, the product was imaged overnight to test the stability of the construct over time.

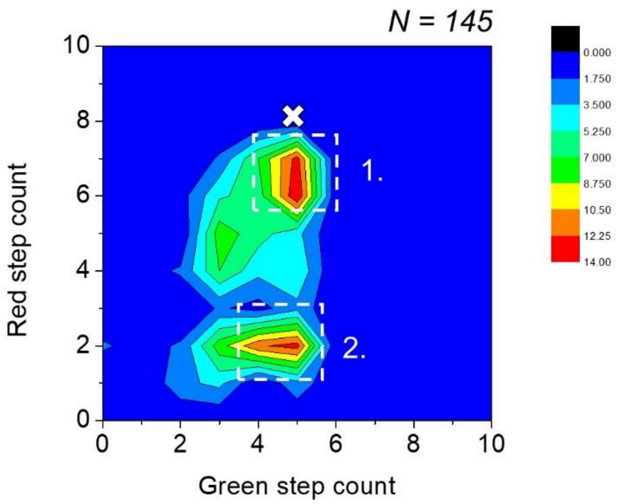


**Figure S14. Construct resilience following incubation overnight.** The two-color correlation analysis for structures fabricated over two days is shown. Desired product is (1.) with ATTO647N-labelled foundation rung, 5 Cy3-labelled rungs, and 5 ATTO647N-labelled rungs. The synthesis was stopped after the first five rungs, as shown in (2.) and then restarted after leaving the surface-grafted material in the fridge overnight. As shown in the two-color plot, equal populations of the structurally complete and the truncated version (the one left overnight) are present, suggesting that fraying/degradation occurred on the last rung of the partially assembled (5 Cy3 bearing rungs) overnight, leading to a subpopulation (roughly 50%) on top of which no additional units may be added the following day.

### *9. Step-count screening algorithm*

We developed a step detection algorithm to automatically estimate the subunit stoichiometry of the synthesized DNA nanotubes from a fluorescence vs. time trace. The step detection algorithm operates through iterative binary clustering of the fluorescence versus time trace (Figure S15). We took care to design an algorithm of linear time complexity, such that the algorithm runtime remains robust to increases in the sampling rate of a photodetector, or other increases to the size of the input data. We demonstrate that the screening algorithm is able to estimate the subunit stoichiometry accurately, both in cases of relative homogeneity and heterogeneity of subunit stoichiometry within a given collection of traces (Figure S17).

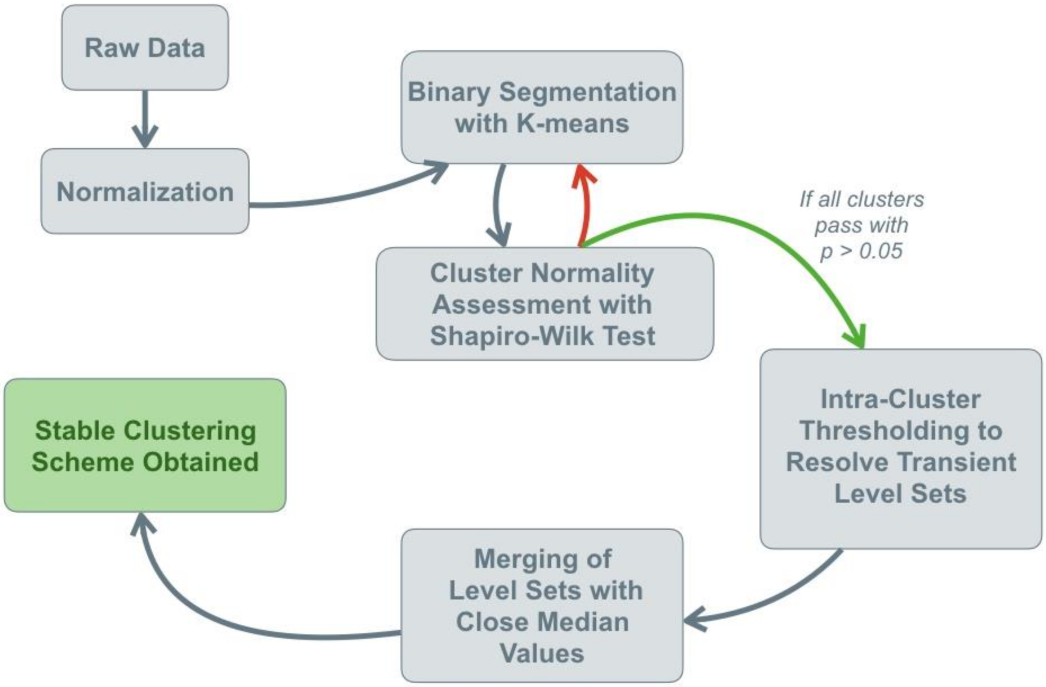


**Figure S15.** Summary of the step detection algorithm.

The screening algorithm operates by performing iterative binary segmentation of the fluorescence vs. time trace. It follows the steps outlined below:
**1)** The fluorescence and time values of the trace are normalized, such that the maximum value for both is 1. This is done to weight the time and fluorescence dimensions equally in the Euclidian distances calculated by the K-means clustering algorithm in subsequent steps.[6]
**2)** The trace is then clustered with the K-means clustering algorithm, constrained to K = 2 clusters.

**3)** The probability distributions of the fluorescence values of each of the resultant clusters is tested against a normal distribution with the Shapiro-Wilk test for normality,[7] with the null hypothesis being that the fluorescence values sample a normal distribution. If, for a given cluster, the null hypothesis is rejected ($p < 0.05$), the cluster is flagged for re-clustering in a subsequent round of K-means clustering.
**4)** Those clusters marked for re-clustering are then clustered again using the K-means algorithm, again constrained to K = 2 clusters, and each resultant cluster is subjected to the Shapiro-Wilk test once again. This process is repeated until all clusters pass the Shapiro-Wilk test with $p > 0.05$, or until a user-defined number of binary segmentations has taken place (typically $\log_2(n_{max} + 1)$), where $n_{max}$ is an estimate of the largest expected subunit stoichiometry given a set of synthetic conditions.
**5)** Following the binary segmentation protocol, all clusters are post-processed (Figure S8). The first post-processing step involves examining each individual cluster for points in the first or last 10% of the range of time values encapsulated in the cluster for points with a deviation of greater than 3σ from the median value of the cluster. This allows for resolution of photobleaching events with very small time separation.
**6)** The second post-processing step involves merging clusters whose median values fall within a user-defined "merging sensitivity" threshold of one another. For the data presented in this paper, we have found that a value between 0.05-0.15 usually results in satisfactory assessments of subunit stoichiometry, though manual examination of clusterings at various values of this parameter is sometimes required to determine the optimal merging sensitivity for a given population of nanotubes under a given set of imaging conditions.
**7)** The fluorescence-time trace is then transformed back to its original magnitude in the fluorescence and time dimensions, and returned to the user with a third, "cluster number" variable, indicating into which cluster each fluorescence-time tuple falls. The subunit stoichiometry is assessed by subtracting one from the total number of clusters.

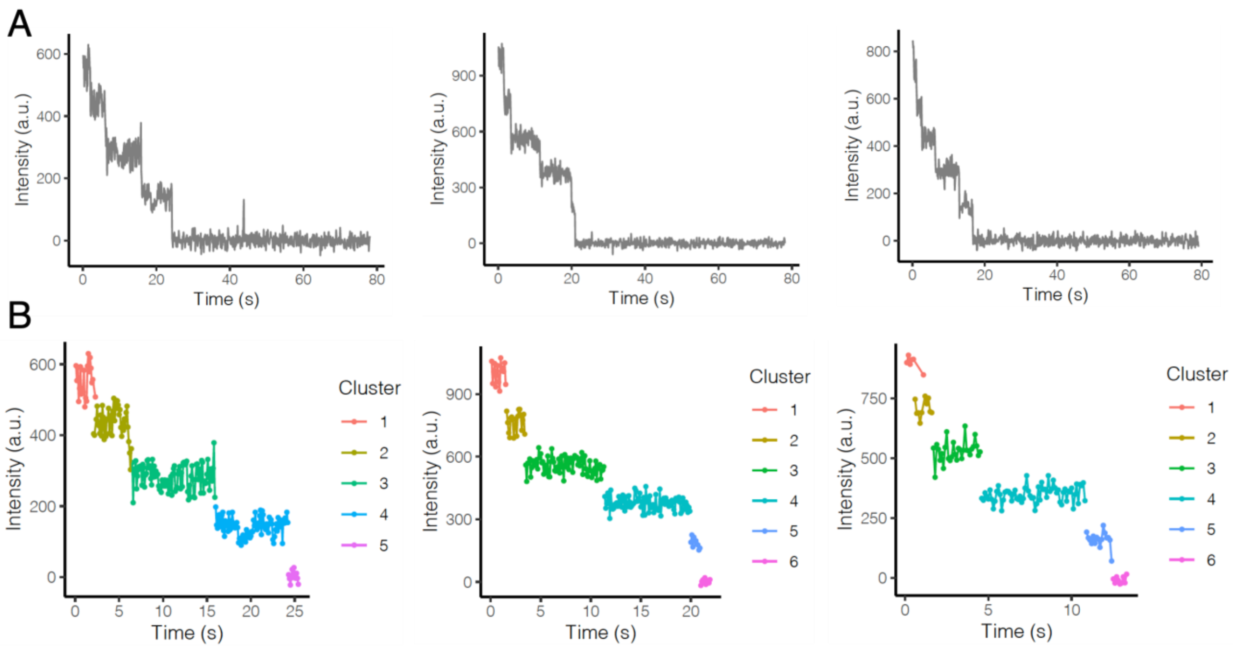

**Figure S16.** (A) Sample data for raw intensity-time trajectories and (B) corresponding algorithm clustering outputs for a 5-rung Cy3-labelled nanotube. The correctly formed product should have 5 fluorophores and therefore 6 identifiable intensity clusters, with the final cluster being the fully bleached product.

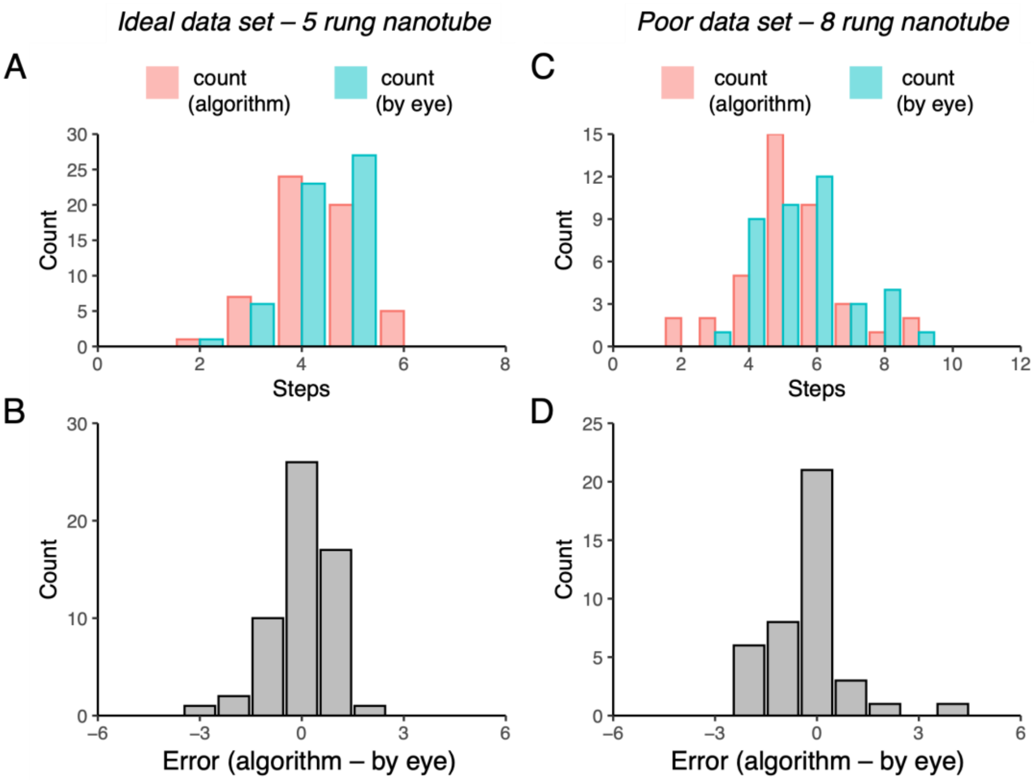


**Figure S17.** The step detection algorithm designed for screening of DNA nanotubes is able to estimate the subunit stoichiometry of a population of nanotubes with a relatively homogenous distribution of subunits (A, B) and a relatively heterogenous distribution of subunits (C, D), enabling the rapid determination of the success of the synthetic protocol without manual analysis of each single molecule trajectory for each nanotube in a given field of view.

#### *9.1 Simulating Intensity-Time Profiles*

To thoroughly characterize the response of Binary K-means to increasing input size, we simulated intensity-time profiles containing stochastic photobleaching events as follows. We defined arbitrarily the probability of photobleaching, $P$, and the de-noised step magnitude, $C$. For a signal of $n$ data points, we generated a list of $n$ random numbers between 0 and 1 from a uniform probability distribution. Every position in the list that contained a number less than $P$ was designated as a step location by changing its value to 1. The values of all other numbers in the list were set to 0.

Then, the simulated trace was created by constructing another list element-by-element, beginning with a starting intensity value $I$. Every time a step location was encountered in the first list, the value $I$ was updated to the value $I$ - $C$, where $C$ was the pre-defined step magnitude. The signal was constructed until List 2 was of length $n$. Finally, Gaussian noise of mean 0 and standard deviation $\sqrt{I}$ was added to each plateau between steps, with $I$ representing the mean intensity between two given steps. We increased *n* far beyond the regime of input size typically encountered in photobleaching experiments presented in this paper.

#### *Time Complexity*

Since technical improvements in photodetector acquisition speeds will invariably lead to fluorescent microscopes with increased time-sampling rates, we explicitly demonstrated the predicted linear time complexity of Binary K-means using simulated data. The runtimes are evaluated on a standard desktop computer, with an Intel i7-4700k CPU.

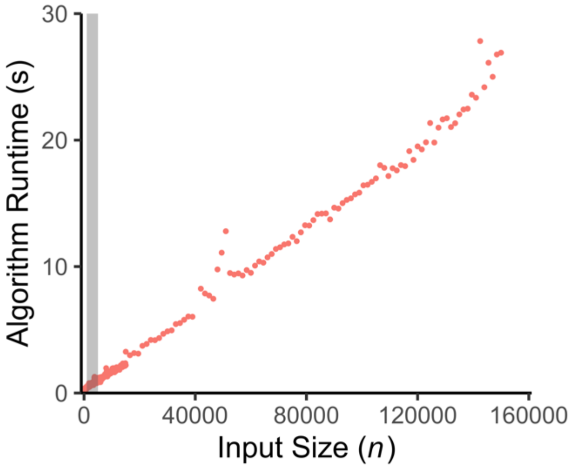


**Figure S18.** Binary K-means is of linear time complexity and very fast in the regime of typical input size, (highlighted in gray). Binary K-means will therefore scale well to the processing of high-resolution intensity-time profiles. The runtimes are measured using a standard desktop computer, with an Intel i7-4700k CPU.

### *10. References*


1. Gidi, Y.; Bayram, S.; Ablenas, C. J.; Blum, A. S.; Cosa, G., Efficient One-Step PEG-Silane Passivation of Glass Surfaces for Single-Molecule Fluorescence Studies. *ACS Appl. Mater. Interfaces* **2018,** *10* (46), 39505-39511.
2. Myong, S.; Bruno, M. M.; Pyle, A. M.; Ha, T., Spring-Loaded Mechanism of DNA Unwinding by Hepatitis C Virus NS3 Helicase. *Science* **2007,** *317* (5837), 513-516.
3. Hariri, A. A.; Hamblin, G. D.; Hardwick, J. S.; Godin, R.; Desjardins, J.-F.; Wiseman, P. W.; Sleiman, H. F.; Cosa, G., Stoichiometry and Dispersity of DNA Nanostructures Using Photobleaching Pair-Correlation Analysis. *Bioconj. Chem.* **2017,** *28* (9), 2340-2349.
4. Cordes, T.; Vogelsang, J.; Tinnefeld, P., On the Mechanism of Trolox as Antiblinking and Antibleaching Reagent. *J. Am. Chem. Soc.* **2009,** *131* (14), 5018-5019.
5. Glembockyte, V.; Cosa, G., Redox-Based Photostabilizing Agents in Fluorescence Imaging: The Hidden Role of Intersystem Crossing in Geminate Radical Ion Pairs. *J. Am. Chem. Soc.* **2017,** *139* (37), 13227-13233.

6. Forgy, E. W., Cluster analysis of multivariate data: efficiency versus interpretability of classifications. *Biometrics* **1965,** *21*, 768-769.
7. Shapiro, S. S.; Wilk, M. B., An analysis of variance test for normality (complete samples). *Biometrika* **1965,** *52* (3/4), 591-611.
8. Cussler, E. L.; Cussler, E. L., *Diffusion: mass transfer in fluid systems*. Cambridge University Press: 2009.
